# The temperature-regulated DEAD-box RNA helicase CrhR interactome: Autoregulation and photosynthesis-related transcripts

**DOI:** 10.1101/2021.03.26.437152

**Authors:** Anzhela Migur, Florian Heyl, Janina Fuss, Afshan Srikumar, Bruno Huettel, Claudia Steglich, Jogadhenu S. S. Prakash, Richard Reinhardt, Rolf Backofen, George W. Owttrim, Wolfgang R. Hess

**Affiliations:** Faculty of Biology, University of Freiburg, Schänzlestr. 1, D-79104 Freiburg, Germany; Department of Computer Science, University of Freiburg, Georges-Koehler-Allee 106, D-79110 Freiburg, Germany; Max Planck-Genome-Centre Cologne, Carl-von-Linné-Weg 10, D-50829 Köln, Germany; Department of Biotechnology & Bioinformatics, School of Life Sciences, University of Hyderabad, Hyderabad, India; Department of Biological Sciences, University of Alberta, Edmonton, Alberta, Canada T6G 2E9

**Keywords:** chloroplasts, CrhR RNA helicase, cyanobacteria, gene expression regulation, photosynthesis, small regulatory RNA, redox regulation, RNA:RNA interaction

## Abstract

RNA helicases play crucial functions in RNA biology. In plants, RNA helicases are encoded by large gene families, performing roles in abiotic stress responses, development, the post-transcriptional regulation of gene expression as well as house-keeping functions. Several of these RNA helicases are targeted to the organelles, mitochondria and chloroplasts. Cyanobacteria are the direct evolutionary ancestors of plant chloroplasts. The cyanobacterium *Synechocystis* 6803 encodes a single DEAD-box RNA helicase, CrhR, that is induced by a range of abiotic stresses, including low temperature. Though the Δ*crhR* mutant exhibits a severe cold-sensitive phenotype, the physiological function(s) performed by CrhR have not been described. To identify transcripts interacting with CrhR, we performed RNA co-immunoprecipitation with extracts from a *Synechocystis crhR* deletion mutant expressing the FLAG-tagged native CrhR or a K57A mutated version with an anticipated enhanced RNA binding. The composition of the interactome was strikingly biased towards photosynthesis-associated and redox-controlled transcripts. A transcript highly enriched in all experiments was the *crhR* mRNA, suggesting an auto-regulatory molecular mechanism. The identified interactome explains the described physiological role of CrhR in response to the redox poise of the photosynthetic electron transport chain and characterizes CrhR as an enzyme with a diverse range of transcripts as molecular targets.

**Highlight:** The cyanobacterial DEAD-box RNA helicase CrhR binds mainly photosynthesis-associated and redox-controlled transcripts connecting its regulation, localization and phenotypes of mutants for the first time with a set of potential RNA targets.

## Introduction

### DEAD-box RNA helicases

The synthesis, maturation, modification and decay of RNA molecules and their interaction with each other and with other cellular components is central to the molecular basis of life. The largest family of enzymes involved in the metabolism of RNA molecules are enzymes belonging to the Superfamily I (SF1) and II (SF2) RNA helicases (Bourgeois *et al*., 2016). DEAD-box RNA helicases, named after the conserved DEAD (Asp-Glu-Ala-Asp) amino acid motif in their core motif, form the largest and most complex group of SF2 RNA helicases (Jarmoskaite and Russell, 2011). DEAD-box RNA helicases are RNA-dependent ATPases (Rocak and Linder, 2004) characterized by the presence of twelve highly conserved motifs (Redder *et al*., 2015). These motifs form the motor core of the helicase, which binds specifically adenine nucleotides and single-stranded RNA in a sequence-independent manner. The main function of these proteins is the conformational rearrangement of RNA by unwinding short double stranded regions. The unwinding reaction can be performed within one RNA molecule or between two duplex-forming RNAs. Besides the unwinding reaction, some RNA helicases are able to anneal single stranded RNAs (Chamot *et al*., 2005; Yang and Jankowsky, 2005). In addition, some RNA helicases act as RNA clamps or facilitate RNA-protein complex dissociation without duplex unwinding (Jankowsky *et al*., 2001; Fairman *et al*., 2004).

### Abiotic stresses and DEAD-box RNA helicases in plants and cyanobacteria

Cold stress, one of the most common stress conditions in nature, frequently induces the expression or activity of DEAD-box RNA helicases. At low temperature, RNA secondary structures are thermodynamically stabilized, which may interfere with their function. RNA helicases can rescue the RNA’s functions by rearranging its secondary structures (Jones *et al*., 1996). This role makes certain RNA helicases more relevant or even conditionally essential at low temperature. For instance, deletion of *csdA* or *srmB* in *Escherichia coli* leads to a cold-sensitive phenotype (Redder *et al*., 2015). If all four DEAD-box RNA helicase genes are deleted, *Bacillus subtilis* is not viable at low temperature (16°C), although the strain grows well at 37°C (Lehnik-Habrink *et al*., 2013).

Most organisms encode several DEAD-box RNA helicases possessing non-complementary roles associated with a variety of physiological functions. Common functions in all organisms involve ribosome biogenesis, RNA turnover and translation, and responses to multiple stress conditions (Py *et al*., 1996; de la Cruz *et al*., 1999; Schneider and Schwer, 2001; Rogers *et al*., 2002; Macovei *et al*., 2012). Plant genomes typically possess larger and more diverse RNA helicase gene families than observed in other systems (Linder and Owttrim, 2009). Hence, their relevance in RNA secondary structure rearrangement under different environmental conditions or in plant development presents interesting lines for investigation. For example, *Arabidopsis thaliana* encodes 58 DEAD-box RNA helicases (Boudet *et al*., 2001), many of which are essential as they are not functionally complementary (Mingam *et al*., 2004). RNA helicases in plants fulfil roles in the defense against viruses (Wu and Nagy, 2020), abiotic stress responses to high salinity (Capel *et al*., 2020), low temperature (Lu *et al*., 2020; Wang *et al*., 2020), and in development (Gong *et al*., 2005). Interestingly, although not complementary, some plant helicases are involved in the same processes, suggesting multiple, independent RNA structure rearrangements are associated with a single physiological response (Huang *et al*., 2016). In addition, several RNA helicases are targeted to the organelles, mitochondria and chloroplasts (Matthes *et al*., 2007; Nawaz and Kang, 2017; Nawaz *et al*., 2018). However, the expanded DEAD-box RNA helicase families make the precise functional characterization of RNA helicases challenging in plants.

Cyanobacteria are the direct evolutionary ancestors of plant chloroplasts (Mereschkowsky, 1905; Margulis, 1981; Martin and Kowallik, 1999; Ponce-Toledo *et al*., 2017). The endosymbiosis of a cyanobacterium not only led to the chloroplast but also had pivotal impact on the composition of the plant nuclear genome (Martin *et al*., 2002). Therefore, analysis of gene function in cyanobacteria is also informative at the higher plant level.

### The cyanobacterium *Synechocystis* encodes a single DEAD-box RNA helicase, CrhR

The unicellular cyanobacterium *Synechocystis* sp. PCC 6803 (from here: *Synechocystis*) encodes the DEAD-box RNA helicase CrhR, for <u>c</u>yanobacterial <u>R</u>NA <u>h</u>elicase redox (Rosana *et al*., 2012a). As defined by a 50 amino acids sequence motif, CrhR is the archetype protein of a new clade within the DEAD-box RNA helicase family (Whitford *et al*., 2021). In contrast to the situation in plants and also in many other bacteria, CrhR (*crhR/slr0083*) is the only DEAD-box RNA helicase encoded in *Synechocystis* (Redder *et al*., 2015). Functionally, CrhR can therefore be expected to be of particular importance as it likely performs multiple functions that from an evolutionary perspective could have been distributed to different members of the complex family of RNA helicases that we observe in plants today (Kiefer *et al*., 2020). *crhR (slr0083)* was originally characterized as a salt and cold-shock inducible protein (Vinnemeier and Hagemann, 1999; Kujat and Owttrim, 2000). Cold stress-inducible RNA helicases were also investigated in the cyanobacterium *Anabaena* sp. PCC 7120 (Chamot *et al*., 1999) and *Synechococcus* sp. WH 7803 (Gierga *et al*., 2012). Detailed analysis indicated that a low temperature stabilization of both transcript and protein contribute to the observed low temperature induction of CrhR (Rosana *et al*., 2012a) and that a *crhR* inactivation mutant was severely impaired morphologically and physiologically at lower but not at higher temperatures (Rosana *et al*., 2012b). In addition to low temperature, *crhR* is induced by a range of abiotic stresses that reduce the electron transport chain, independent of temperature shift (Vinnemeier and Hagemann, 1999; Kujat and Owttrim, 2000; Ritter *et al*., 2020). The gene *crhR* (*slr0083*) is located in a dicistronic operon together with *rimO* (*slr0082*), whose putative protein product has 38% identity with RimO, a ribosomal protein S12 methylthiotransferase (UniProtKB P0AEI4), and 29% identity with the paralogous tRNA methylthiolase MiaB (Uni-ProtKB P0AEI1) from *E. coli* (Rosana *et al*., 2020). The auto-regulated enhanced operon discoordination and processing of the *crhR* mRNA from the dicistronic operon RNA adds further complexity to its regulation (Rosana *et al*., 2012; Rosana *et al*., 2020). In addition, upon temperature upshift CrhR undergoes rapid repression via conditional proteolysis at the post-translational level through an unknown mechanism (Tarassova *et al*., 2014). In localization studies, CrhR localized to the thylakoid membranes and co-sedimented with degradosome and polysome complexes (Rosana *et al*., 2016), consistent with the recent co-fractionation analysis using Grad-seq (Riediger *et al*., 2021).

Although the molecular effects of *crhR* deletion or inactivation were studied at both the transcriptome (Prakash *et al*., 2010; Georg *et al*., 2019) and proteome level (Rowland *et al*., 2011), the direct RNA targets of CrhR and interacting protein partners have not been identified. To monitor transcriptome-wide binding of CrhR, we immuno-precipitated the RNA species interacting with a FLAG-tagged version of the native RNA helicase expressed in a Δ*crhR* background (subsequently called CrhR_WT_). We performed UV cross-linking *in vivo* with CrhR_WT_ cultures grown at the standard growth temperature of 30°C or exposed to 20°C for 2 h (low temperature stress). A possible obstacle in the analysis of RNA helicases can be their transient interaction with RNA molecules followed by rapid ATP-dependent dissociation (Linder and Jankowsky, 2011). Therefore, we introduced a K57A mutation located in the predicted Walker A ATP-binding motif I (GTGKT) that is conserved in CrhR (Tanner and Linder, 2001). Mutation of the conserved lysine or the last threonine in this motif is known to interfere with the ATPase activity of DEAD-box RNA helicases (Cordin *et al*., 2006). Plasmid encoded K57A was conjugated into the Δ *crhR* background yielding strain CrhR_K57A_. This strain was used for comparison applying UV cross-linking *in vivo* at 30°C followed by co-immunoprecipitation (co-IP) and RNA sequence analysis as for the CrhR_WT_ strain.

Altogether, 119 RNA sequences were significantly enriched in at least one experiment. Functional analysis of the specifically enriched RNAs indicated a striking preference for CrhR interaction with transcripts associated with photosynthesis but also transcripts associated with RNA metabolism, among them *crhR* itself. The enrichment of *crhR*– transcripts with the tagged CrhR protein is consistent with an auto-regulatory mechanism in which CrhR controls its own expression at the post-transcriptional level. Broader implications were revealed by the potential RNA targets having previously been characterized as controlled by the transcription factor RpaB (Riediger *et al*., 2019), which is of interest in conjunction with the known ETC redox poise regulation of *crhR* expression (Kujat and Owttrim, 2000; Ritter *et al*., 2020). Overall, the results connect the known association of CrhR with both the thylakoid membrane and ribosomes and its cold-induced and redox poise-controlled expression with a specific set of potential RNA targets comprising a subset of the *Synechocystis* transcriptome.

## Materials and methods

### Bacterial strains

The *Synechocystis*Δ*crhR* mutant (Prakash *et al*., 2010) was used for the construction of three different strains. To establish ectopic expression of tagged CrhR, the triple FLAG-tag was introduced at the 5’-end of the wild type *crhR*/*slr0083* gene by PCR in three steps. First, *crhR* was amplified with the primers 3xFL-CrhR-F1 and 3xFL-CrhR-R (see **Table S1** for the sequences of oligonucleotide primers used in this work), re-amplified with primers 3xFL-CrhR-F2 and 3xFL-CrhR-R, followed by another reamplification using the primers 3xFL-CrhR-F3 and 3xFL-CrhR-R. The resulting amplicon 3xFLAG-*crhR* was digested with *Nde*I and *Xba*I and ligated into pJET1.2. The copper inducible promoter, P_petE_ (Zhang *et al*., 1992) was amplified with the primers petE-FP and petE-RP and cloned into pJET1.2 via Gibson assembly. The *rrnB* terminator was amplified from pBAD using primers rrnB-TT-F and rrnB-TT-R and ligated into pJET1.2 via *Sac*I and *Xba*I restriction sites yielding pJET1.2::P*_petE_*::3xFLAG-*crhR*::*rrnB*T.

To introduce the K57A mutation into *crhR*, primers crhR(K57A)_inverse_fw and crhR(K57A)_inverse_rv were used in an inverse PCR to generate CrhR_K57A_. Both the pJET1.2::P*_petE_*::3xFLAG-*crhR*::*rrnB*T and CrhR_K57A_.constructs contain the P*_petE_* promoter followed by the triple FLAG tag translationally fused to the *crhR* ORF followed by the *rrnB* terminator. To eliminate the *crhR* gene and create the negative control Δ*crhR*_control_, the vector pJET1.2::P*petE*::3xFLAG-*crhR*::*rrnB*T was inverse PCR amplified with primers petE-3xF-rrnb_neg_ctrl_FP and petE-3xF-rrnb_neg_ctrl_RP carrying *Sac*I restriction sites on the 5′ overhangs, cleaved with *Sac*I and self-ligated.

For cloning of the constructs into pVZ321 (Zinchenko *et al*., 1999), the constructs were amplified from pJET1.2 using the primers petE_foR-pVZ_FP and rrnB_foR-pVZ_RP carrying *Eco*RI restriction sites. The amplified constructs were inserted into the *Eco*RI restriction site of pVZ321. This insertion resulted in disruption of the chloramphenicol resistance cassette of pVZ321, so that the final constructs pVZ321::P*_petE_*::3xFLAG-*crhR*::*rrnB*T (strain CrhR_WT_), pVZ321::P*_petE_*::3xFLAG-*crhR*(K57A)::*rrnB*T (strain CrhR_K57A_) and pVZ321::P*_petE_*::3xFLAG::*rrnB*T (strain Δ*crhR*_control_) possessed only kanamycin resistance. The resulting pVZ321 plasmids were transferred into the Δ*crhR Synechocystis* strain (Prakash *et al*., 2010) via triparental mating with *E. coli* J53/RP4 and TOP10F’ (Scholz *et al*., 2013). Exconjugants were selected on agar plates containing 20 μg/ml spectinomycin (Sp) and 50 μg/ml kanamycin (Km) in BG-11 (Rippka *et al*., 1979).

### Culture conditions

*E. coli* strains were grown in liquid LB medium (10 g l^−1^ bacto-tryptone, 5 g l^−1^ bacto-yeast extract, 10 g l^−1^ NaCl), with continuous agitation or on agar-solidified (1.5% [w/v] bacto agar) LB, supplemented with appropriate antibiotics, at 37°C.

*Synechocystis* strains were cultivated in the presence of appropriate antibiotics either in Erlenmeyer flasks in BG-11 medium (Rippka *et al*., 1979) or in 100 ml two-tier vessel CellDEG cultivators (CellDEG GmbH (Bähr *et al*., 2016)) at the indicated temperatures in fresh water organisms (FWO) medium with shaking. FWO medium consists of 50 mM NaNO_3_, 15 mM KNO_3_, 2 mM MgSO_4_, 0.5 mM CaCl_2_, 0.025 mM H_3_BO_3_, 0.15 mM FeCl_3_/Na_2_EDTA, 1.6 mM KH_2_PO_4_, 2.4 mM K_2_HPO_4_, 10 mM NaHCO_3_, 10 nM MnCl_2_, 1 nM ZnSO_4_, 2 nM Na_2_MoO_4_, 2 nM CuSO_4_, 30 pM CoCl_2_. High CO_2_ concentrations were delivered to the flasks from a carbonate buffer (1.447 M K_2_CO_3_ and 12.486 M KHCO_3_) through a highly gas-permeable polypropylene membrane. The CellDEG cultivators were inoculated at an OD_750_ ∼0.8 and a light intensity of 150 μmol photons m^−2^s^−1^. Light intensity was adjusted to culture density; therefore, 24 h after the start of inoculation light intensity was increased to 300 μmol photons m^−2^s^−1^ and to 500 μmol photons m^−2^ s^−1^ after another 24 h. To induce gene expression from the Cu^2+^-responsive promoter P*_petE_* in the CellDEG system, 6 μM CuSO_4_ was added to the medium. The cells were harvested at OD_750_ ∼20, which was reached upon 72 hours of cultivation.

### FLAG-tag affinity purification, UV crosslinking and RNA co-IP

*Synechocystis* cultures were grown in CellDEG cultivators to an OD_750_ ∼20 at 30°C. For cold stress, cultures were exposed to 20°C for 2 h. The cells were placed in a 20 x 20 cm Petri dish and irradiated three times on ice with 500 mJ/cm^−2^ UV-C light (254 nm) in a Stratalinker® 2400 for a total irradiation time of 3 min. The treated cells were harvested by centrifugation (4,000 x g, 10 min, 4°C). Cell pellets were resuspended in precooled TBS buffer (50 mM Tris-HCl (pH 7.5)), containing cOmplete™ Protease Inhibitor Cocktail (Roche), washed once with TBS and centrifuged again. The washed and cross-linked cells were suspended in TBS buffer and 0.5 volume of glass beads were added. Cells were disrupted mechanically applying 5 cycles at 6,500 rpm with 10 s breaks on ice between the cycles in a Precellys® 24 homogenizer. The cell lysate was separated by centrifugation (13,000 x g, 15 min, 4°C) into two fractions, one containing membrane proteins and the other containing soluble proteins. The latter was incubated with 50 μl of Anti-FLAG® M2 magnetic beads (Sigma Aldrich) to allow binding of FLAG-tagged proteins at 4°C for 4 hours. Subsequently, beads were washed twice in TBS-containing cOmplete™ Protease Inhibitor Cocktail followed by a wash step that removed the non-crosslinked RNA by incubation in 500 μl RNA elution buffer (25 mM Tris-HCl (pH7.5), 2 M NaCl) for 15 min at room temperature. The crosslinked RNA was released by digestion of the FLAG-tagged protein with 20 μg of proteinase K for 10 min at room temperature, purified using the RNA Clean & Concentrator-5 Kit (Zymo Research) and utilized for the generation of cDNA libraries for Illumina sequencing as described below. All co-immunoprecipitation experiments were performed in biological duplicates.

### RNA preparation

*Synechocystis* cell pellets were disrupted by incubation for 15 min at 65 °C in PGTX as described by Pinto *et al*. (2009). Chloroform/isoamyl alcohol (IAA) (24:1) extraction was performed for 10 min at room temperature followed by centrifugation at 3,270 x g for 15 min at room temperature for phase separation. The chloroform/IAA extraction was repeated on the aqueous phase, followed by isopropanol precipitation. Precipitated RNA was pelleted by centrifugation (13,000 x g, 30 min, 4°C), washed with 70% ethanol, air-dried and resuspended in nuclease-free water. RNA concentration was determined on a NanoDrop ND-1000 spectrophotometer. RNA purity and quality was evaluated on a Fragment Analyzer parallel capillary electrophoresis system (Agilent).

### Preparation of cDNA libraries

To initiate cDNA library preparation, 0.3-5 µg of total RNA was treated with 4 U of TURBO DNase (Life Technologies) in 2 consecutive incubation steps, each at 37°C for 15 min as indicated by the manufacturer. Next, RNA clean-up and separation into small (17-200 nt) and large (>200 nt) RNA fractions was performed using RNA Clean & Concentrator columns (Zymo Research). The large RNA fraction was fragmented according to the protocol by Pfeifer-Sancar *et al*. (2013). After RNA Clean & Concentrator column purification, both fractions were combined and all subsequent steps performed as previously described (Pfeifer-Sancar *et al*., 2013) except that the clean-up of RNA 5’-pholyphosphatase (Epicentre)-treated samples was performed by Clean & Concentrator column purification. For RNA adapter ligation, the UGA linker (Supplementary **Table S1**) was used. After RNA adapter ligation and cDNA synthesis, the samples were excised from 2% agarose gels and purified using the NucleoSpin Gel and PCR Clean-up kit (Macherey-Nagel) using NTC buffer for solubilization of the gel slices. After PCR amplification, residual primers were removed by adding 10 µl of ExoSAP-IT (USB) to 50 µl of PCR, and with samples incubated for 15 min at 37°C followed by heat inactivation of the enzyme for 15 min at 85°C. The samples were cleaned-up using a NucleoSpin Gel and PCR Clean-up kit. The quality of RNA and DNA was analyzed on a Fragment Analyzer parallel capillary electrophoresis system (Agilent). Sequencing of libraries was performed on a HiSeq 3000 sequencer with a 2 x 150 bp read mode.

### Bioinformatics analyses

Analysis of the raw reads from the RNA sequencing was performed using the Galaxy web platform (https://usegalaxy.eu/ (Afgan *et al*., 2018)). Illumina paired-end reads were trimmed, adapter sequences and reads shorter than 14 nt filtered out using cutadapt 1.16 (Martin, 2011). The remaining reads were mapped to the chromosome and plasmids of *Synechocystis* using bowtie2 version 2.3.4.3 with the following selected parameters for paired-end reads: -I 0 -X 500 --fr --no-mixed --no-discordant --very-sensitive(Langmead and Salzberg, 2012). Peak calling of the mapped reads was performed by PEAKachu 0.1.0.2 with the parameters: --pairwise_replicates -- norm_method deseq --mad_multiplier 2.0 –fc_cutoff 1 --padj_treshold 0.05. The total number of reads as well as those that mapped to the *Synechocystis* genome are listed in **Supplementary Material, Table S1**.

RNA secondary structures were predicted using the RNAfold WebServer as part of the ViennaRNA Websuite (Gruber *et al*., 2015) with default parameters. Structures were visualized for publication in the VARNA Applet version 3.93 (Darty *et al*., 2009).

### Generation of recombinant CrhR

To obtain recombinant, C-terminal His-tagged CrhR, the *crhR* gene was amplified from the constructs pVZ321::P*petE*::3xFLAG-*crhR*::*rrnB*T with primers pQE70_crhR-fw and pQE70_crhR-rv (**Table S1**). The pQE70 vector was amplified with primers pQE_GIBSOn-fw and pQE_GIBSOn-rv. The fragments were mixed and transformed into *E. coli* Top10 F’ using AQUA cloning (Beyer *et al*., 2015) yielding strain pQE70:crhR-6xHis. For recombinant protein expression, the construct pQE70::*crhR*-6xHis was transformed into *E. coli* M15. Overnight cultures were diluted 1:100 in fresh LB medium supplemented with 50 µg/ml ampicillin, and grown to an OD_600_ of 0.7 at 37°C. Protein expression was induced by adding IPTG to 1□mM final concentration. Three hours after induction, cells were harvested by centrifugation at 6,000 x g for 10 min at room temperature. Cell pellets were resuspended in lysis buffer (50□mM NaH_2_PO_4_ (pH 8), 1□M NaCl, 10% glycerol, 15□mM imidazole, cOmplete™ Protease Inhibitor Cocktail (Roche)) and lysed using the One Shot constant cell disruption system (Constant Systems Limited, U.K.) at 2.4□kbar. Cell lysates were cleared by centrifugation at 13,000 g for 30□min at 4°C and the lysate was filtered through 0.45□ µm Supor-450 filters (Pall). Recombinant proteins were immobilized on a HiTrap Talon crude 1 ml column (GE Healthcare), equilibrated with buffer A (50□mM NaH_2_PO_4_ (pH 8), 500□mM NaCl), and eluted with elution buffer B (50□mM NaH_2_PO_4_ (pH 8), 500□mM imidazole, 500□mM NaCl). The protein concentration was calculated with a Bradford assay using bovine serum albumin as the standard.

### Cy3 labelling of RNA

To initiate Cy3 RNA labelling for EMSA analysis, two volumes of 6 mM KIO_4_ were added to 4 μg RNA and incubated at room temperature in the dark for 1 h. The RNA was precipitated at −20°C for 1 h by addition of one volume of ethylene glycol/H_2_O (1:1), 2.9 volumes of 100% ethanol, and 0.1 volume of 3.3 M NaCl and pelleted at 13,000 x g for 30 min at 4°C. The RNA pellet was washed with 70 % ethanol, air dried, and resuspended in 10 μl of 50 mM Cy3 dye dissolved in DMSO (Thermo Fisher Scientific) and incubated at 37°C in the dark for 2 h. Two volumes of 0.1 M Tris-HCl (pH 7.5) were added followed by the addition of 20 mM freshly prepared NaBH_4_ and the reaction mix incubated at 4°C in the dark for 30 min. The reaction was quenched and RNA precipitated by ethanol precipitation.

### Electrophoretic mobility shift assays

Binding of different amounts of recombinant CrhR (0.1 to 5 pmol) to 0.2 pmol of Cy3 labelled RNA was performed in 20 mM HEPES-KOH (pH 8.3) buffer containing 3 mM MgCl_2_, 1 mM DTT, 500 μg/ml BSA. As a substrate competitor, 1□μg of LightShift Poly (dIdC) (Thermo Fischer Scientific) was included in each assay. The reaction was incubated at room temperature for 15 min prior to separation on 2% agarose-TAE gels. The signals were visualized with a Laser Scanner Typhoon FLA 9500 (GE Healthcare) using a green light laser at a wavelength of 532 nm and a Cy3 filter (LPG, DGR1, BPG1.

## Results

### Physiological consequences of manipulation of CrhR sequence and abundance

To gain insight into the CrhR interactome, we performed immunoprecipitation and isolation of tagged CrhR-RNA complexes from *in vivo* UV cross-linked *Synechocystis* cells. A previously characterized helicase deletion mutant, Δ*crhR*, generated by replacement of the *crhR* gene (*slr0083*) by a spectinomycin resistance gene (Prakash *et al*., 2010), served as the genetic background in which three strains were constructed. One, ectopically expressing the triple FLAG-tagged CrhR (CrhR_WT_), and the second expressing the triple FLAG-tagged CrhR(K57A) mutant (CrhR_K57A_) from the conjugative vector pVZ321 under control of the copper inducible promoter, P_petE_ (Zhang *et al*., 1992) (**Fig. 1A**). The introduced K57A mutation should interfere with ATP binding and ATPase activity of CrhR (Tanner and Linder, 2001) and therefore delay or reduce its capability to dissociate from bound RNA. This mutation is predicted to stabilize CrhR-target transcript interaction, thereby aiding our ability to identify especially low-abundant transcripts. As a control, a third strain was constructed with pVZ321 carrying a short reading frame encoding a triple FLAG tag and identical promoter and terminator sequences as the other two strains (Δ*crhR*_control_). Three experiments were performed with these strains (**Fig. 1B**). The proper regulation and expression of FLAG-tagged CrhR proteins was confirmed by western blotting (**Fig. S1**). We noticed that under identical conditions the expression level of CrhR_K57A_ was higher than that of CrhR_WT_, although (with the exception of the single substitution) both constructs including the promoter, 5’ and 3’ sequences, were identical. Therefore, the protein carrying the K57A substitution appears to be stabilized in the absence of functional CrhR.

**Fig. 1.**
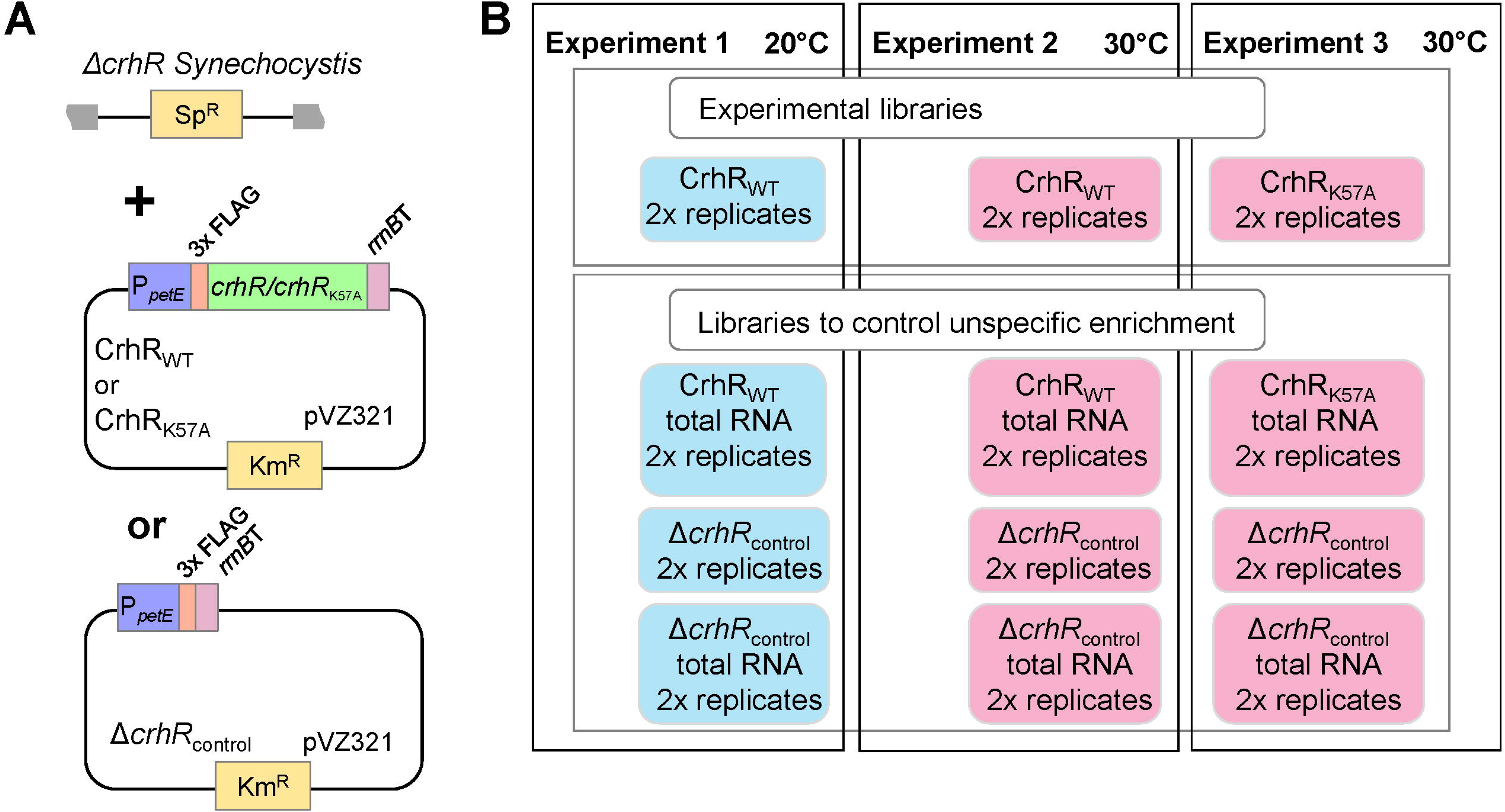
Overview on the experimental flow. **(A)** Schematic representation of the cyanobacterial strains created for the chosen strategy. A deletion mutant, Δ*crhR*, previously generated by replacement of the *crhR* gene (*slr0083*) by a spectinomycin (Sp^R^) resistance gene by homologous recombination (Prakash *et al*., 2010) served as the initial platform. In this background, strains were engineered that express the native form of CrhR (CrhR_WT_) or a K57A substitution (CrhR_K57A_), both translationally fused to an N-terminal triple FLAG tag in the conjugative vector pVZ321 under control of the copper inducible *P_petE_* promoter. A strain expressing only the 3xFLAG-tag was constructed as a control (Δ*crhR*_control_). **(B)** Experimental design. Experiment 1: co-IP of the UV-crosslinked RNA-CrhR_WT_ complexes was performed from *Synechocystis* CrhR_WT_ cultures exposed to cold stress at 20°C for 2 h or, Experiment 2, grown at 30°C. Experiment 3: co-IP of UV-crosslinked RNA-CrhR_K57A_ complexes was performed from cultures grown at 30°C. In parallel, to control for non-specific enrichment, an RNA mock co-IP was performed with the Δ*crhR*_control_ strain. Cultures in Experiments 1 to 3 were grown in FWO medium using the CellDEG system. Each experiment was performed in biological duplicates and cDNA libraries were prepared from the enriched RNA and from the total RNA of all respective cultures omitting the co-IP step.

We followed the growth of the three generated strains and wild type over 25 days at 20°C. Growth of Δ*crhR*_control_ was dramatically reduced compared to wild type cells, while growth of CrhR_WT_ was similar to the unmodified wild type (**Fig. 2A**). Thus, ectopically transcribed, FLAG-tagged CrhR in the CrhR_WT_ strain complemented the *crhR* deletion at this temperature. Interestingly, complementation in strain CrhR_K57A_ led to a partial restoration of the cold-sensitive phenotype, i.e. growth of CrhR_K57A_ was reduced *crhR*_control_ compared to the wild type but not to the same extent as of Δ*crhR*_control_ (**Fig. 2A**). In addition, a strong reduction in pigment content in CrhR_K57A_ and in Δ*crhR*_control_ was apparent compared to the other two strains (**Fig. 2B, C**). These variations were congruent with previously identified changes in light-harvesting pigment composition upon *crhR* inactivation (Rosana *et al*., 2012b). We concluded that the FLAG tag did not interfere with the enzymatic and regulatory functions of CrhR and therefore these lines were suitable for more detailed analyses.

**Fig. 2.**
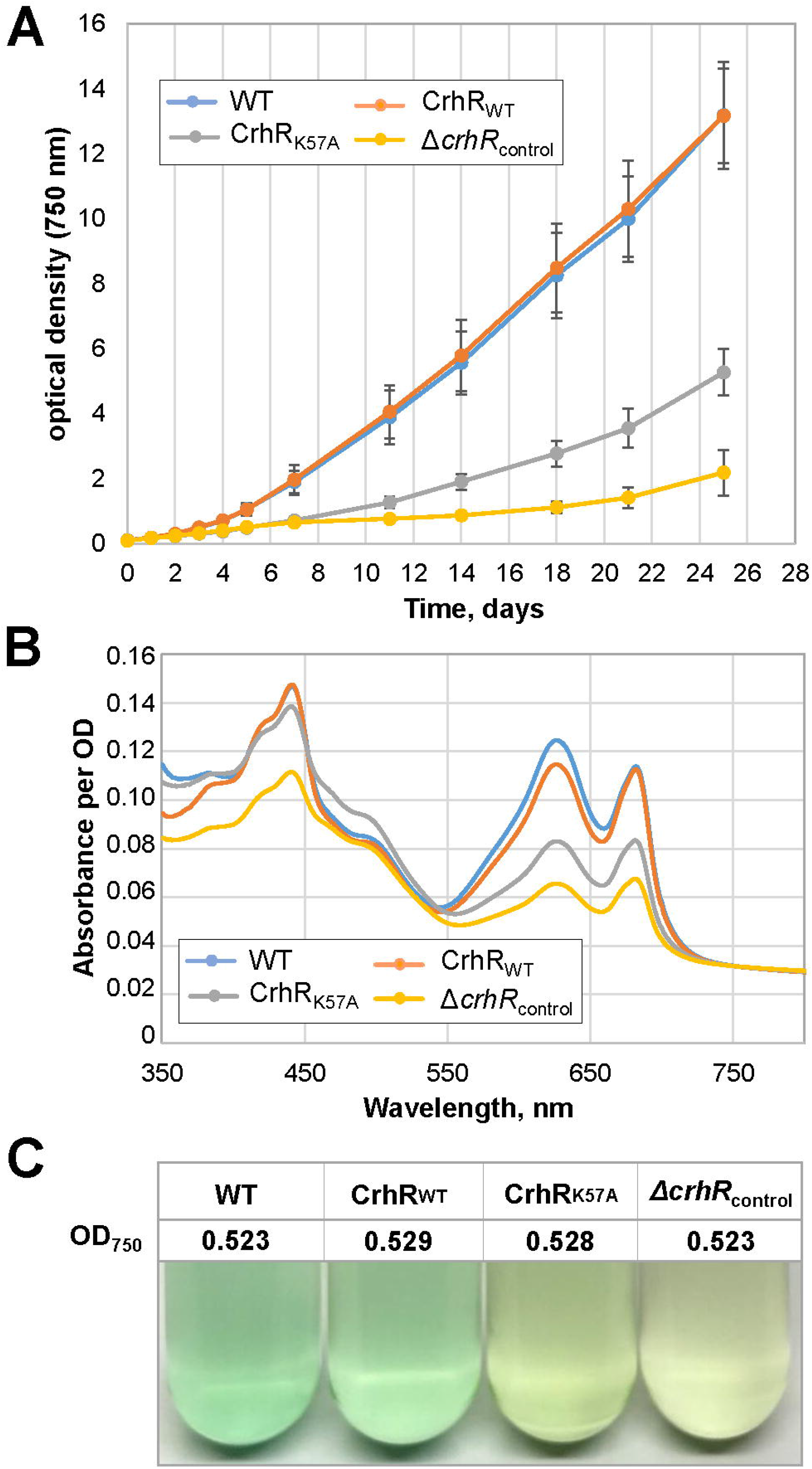
Phenotypiccharacterization of wild type, CrhR_WT_, CrhR_K57A_ and the. Δ***crhR*_control_ strains. (A)** Growth of the wild type (WT), CrhR_WT_, CrhR_K57A_ and Δ*crhR*_control_strains was determined spectrophotometrically by measuring the OD_750_ of cultures grown at 20°C in FWO medium. The experiments were performed in biological triplicates and standard deviations are indicated. **(B)** Pigmentation in the wild type (WT), CrhR_WT_, CrhR_K57A_ and Δ*crhR*_control_ cultures at an OD_750_ of 0.52 grown at 20°C in BG-11 medium. Spectra were normalized to 750 nm. **(C)** Visual appearance of the four strains at the indicated OD_750_. Strains were grown in the presence of antibiotics and 2 μM Cu_2+_.

### RNA co-immunoprecipitations

The RNA helicase CrhR performs unwinding and annealing of RNA (Chamot *et al*., 2005), reactions for which mainly transient interactions with RNA can be assumed including binding to RNA followed by rapid dissociation. This makes the experimental mapping of an RNA helicase interactome more challenging than for an RNA-binding protein. Therefore, we performed covalent crosslinking *in vivo* to enrich RNA-CrhR complexes sufficient for RNA and library preparation. CrhR_WT_ and CrhR_K57A_ cultures grown at the standard growth temperature of 30°C were irradiated with 254 nm UV-C light at 500 mJ/cm^2^ (Holmqvist *et al*., 2016). To identify potentially different interacting partners, the experiment was also performed on CrhR_WT_ cultures exposed to cold stress at 20°C for 2 hours. In parallel, total transcriptome data were generated for all strains and conditions (see overview in **Fig. 1B**). The total and mapped read numbers are listed in **Supplementary Material, Table S2**. CrhR interacting RNAs were predicted with the peak calling algorithm PEAKachu (Holmqvist *et al*., 2016). In total, we obtained 98 different peaks with statistically significant enrichment (padj < 0.05, log_2_FC > 0) in the three experiments corresponding to 64 different genes (**Supplementary Material, Tables S3 to S5**). CrhR interacting RNAs were most abundant in experiment 3 using CrhR_K57A_ as bait (75 peaks), while 28 and 16 peaks were identified in the experiments 1 (20°C) and 2 (30°C) using CrhR_WT_. The higher number of peaks using CrhR_K57A_ as bait matches the assumption that the K57A substitution would stabilize CrhR-RNA interactions. In addition, the somewhat higher expression level of CrhR_K57A_ compared to CrhR_WT_ (**Fig. S1**) may have contributed as well.

Interestingly, the co-IPs frequently identified only specific regions of RNA transcripts which could provide insight into both the molecular mechanism and also the functional roles CrhR is performing. Typical patterns of enriched transcripts are visualized in **Fig. 3**. The predominant detected interactions were associated with segments containing the 5’-untranslated region (5’ UTR), start codon and a few codons into the reading frame (**Fig. 3A**). A related pattern also included the 5’ UTR, start codon and initial codons but actually matched the entire intergenic space between two genes in an operon (**Fig. 3B**). A strikingly different pattern was observed in which multiple sharp peaks indicated a multitude of interactions along the length of the dicistronic operon, as illustrated for the *psaA*-*psaB* operon (**Fig. 3C**). Nevertheless, as observed for other target transcripts, the most pronounced interactions corresponded to the 5’ UTR, start and first codons of *psaA*, the first gene of this operon (**Fig. 3C**). We noticed that the read distribution in the enriched RNA segments resembled somewhat the total RNA-seq read distribution. The reason likely is that the CrhR-mRNA interaction lowers the accessibility for endoribonucleases and thereby protects the respective transcript segments from degradation *in vivo*.

**Fig. 3.**
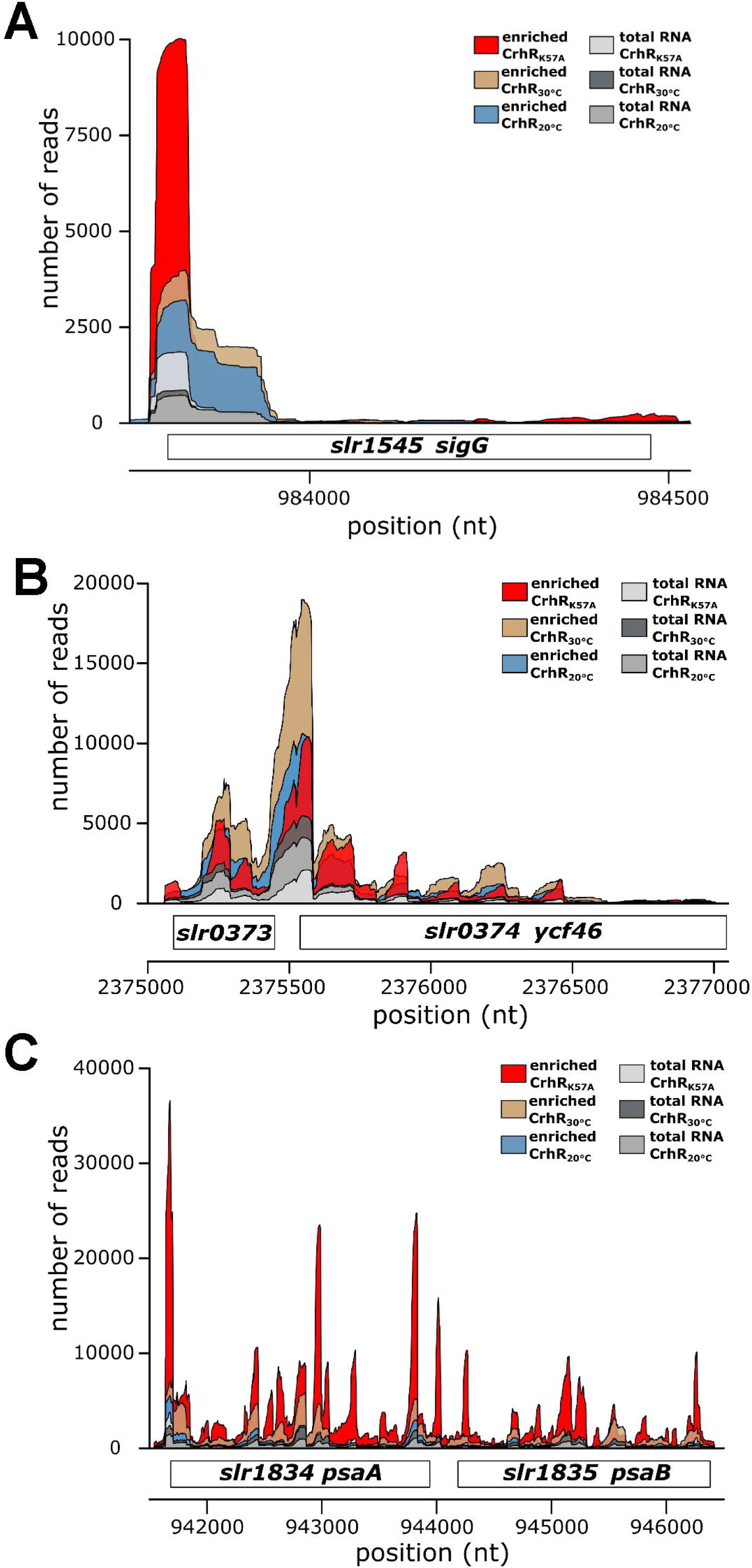
Peak pattern of transcripts enriched in CrhR co-IPs. **(A)** The region containing the 5’ UTR, start codon and following codons were enriched for gene *slr1545* encoding the RNA polymerase sigma factor SigG. This peak was significant in all three experiments, using CrhR_K57A_ or CrhR_WT_ as bait proteins at the standard growth temperature of 30°C (read coverage in red and brown) as well as with CrhR_WT_ at 20°C (cold stress, coverage in blue). **(B)** Reads yielding a peak in the middle of the first gene and extending into the entire intergenic space between two genes and including the 5’ UTR, start codon and following codons of the second gene were recovered for the dicistronic operon *slr0373* and *slr0374*. **(C)** Multiple sharp peaks indicating a multitude of interactions between CrhR and the dicistronic *psaAB* mRNA. *psaAB* encode the PSI P700 apoprotein subunits Ia and A2. In all three panels, the enrichment from reads recovered from CrhR co-IP experiments is shown (red, brown and blue) together with the coverage from RNA-Seq analyses of the respective total transcriptomes in shades of grey as indicated. The numbers along the x-axis correspond to chromosomal nucleotide positions in *Synechocystis* (Genbank accession number NC_000911.1), while the normalized numbers of reads are given on the y-axis.

### Transcripts encoding photosynthesis-associated proteins or with predicted roles in redox regulation are enhanced in the CrhR interactome

Functional analysis over the complete dataset using GO terms (Ashburner *et al*., 2000; Gene Ontology Consortium, 2021) identified “photosynthetic membrane” (q_adj_ = 1.18E-04), “thylakoid” (q_adj_ = 2.74E-04) and “photosystem” (q_adj_ = 2.74E-04) as the dominating cellular components encoded by the enriched transcripts.

Accordingly, detailed analysis indicated that most of the enriched transcripts were associated with photosynthesis (**Table 1;** for the exact coordinates of recovered transcripts, enrichment factors and adjusted p values in each of the experiments, see **Supplementary Material, Tables S3 to S5**). Several of these transcripts encode protein components of the two photosystems, PSI and PSII. PSI proteins included *psaL/slr1655* for PSI subunit XI, *slr1834/psaA* and *slr1835/psaB* encoding the P700 apoprotein subunits Ia and A2 **(Fig. 3C)**, *ssr2831/psaE* encoding the PSI reaction center subunit IV. Transcripts encoding PSII subunits included *slr0906/psbB* for the CP47 reaction center protein, *slr0927/psbD*, *slr1311/psbA2* and *smr0009* for the reaction center proteins D2, D1 and PsbN.

**Table 1.**
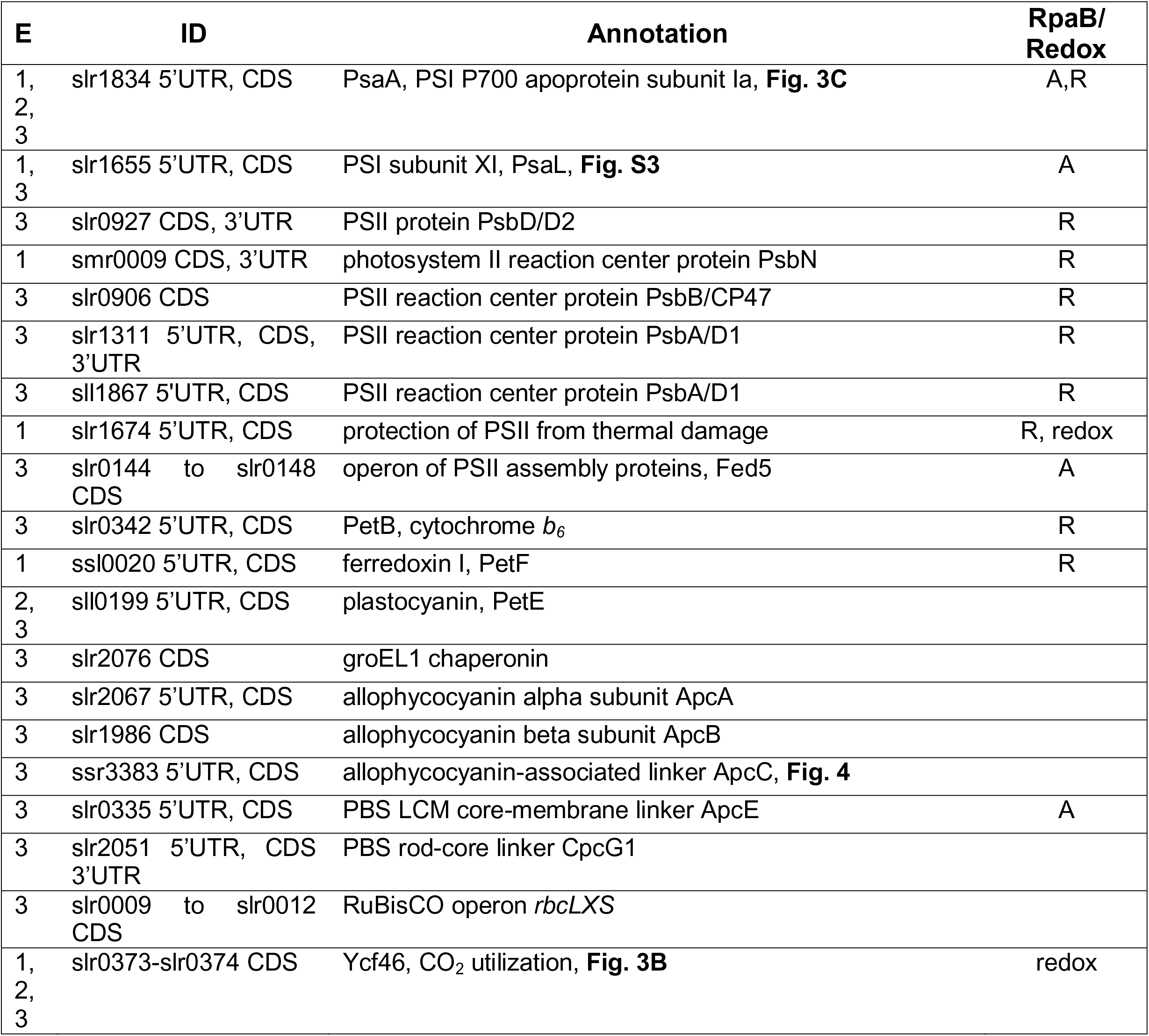
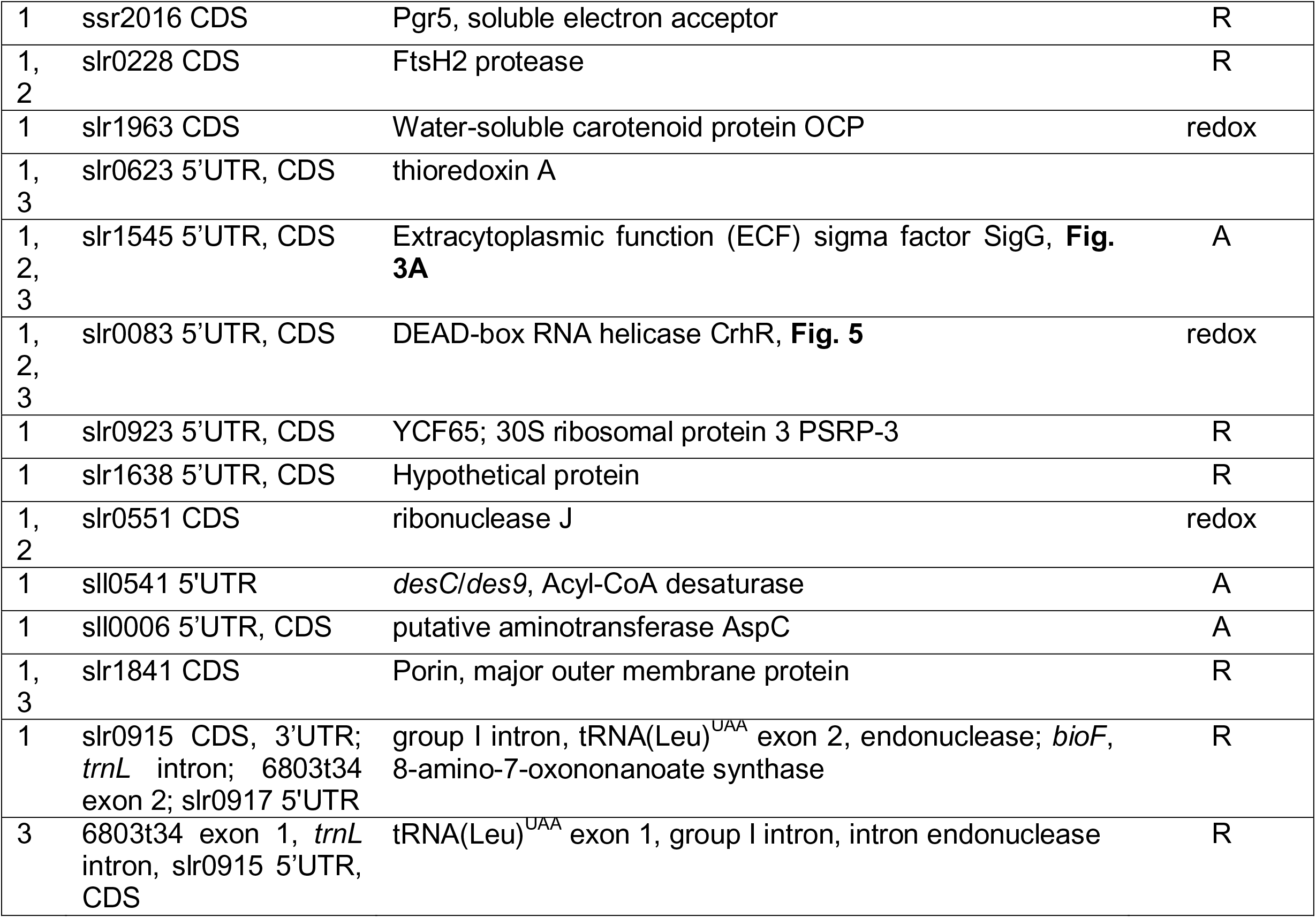
Abundant transcripts recovered in CrhR co-IPs. Assignment to one of the three experiments (**Fig. 1**) is given by the numbers in the first column, followed by gene IDs according to Genbank file NC_000911 and localization of the peak within the coding region (CDS) or UTR. The final two rows provide the annotation of the interacting RNAs and information on their transcriptional regulation, either activation (A) or repression (R) by the transcription factor RpaB under low light (Riediger *et al*., 2019) or with “redox”, if they were classified as redox-responsive genes (Hihara *et al*., 2003; Ritter *et al*., 2020). Detailed information on peak locations, enrichment factors and adjusted p values generated by the PEAKachu algorithm (Holmqvist *et al*., 2016) are given in **Tables S3** to **S5**.

Several enriched transcripts encoding proteins that are not directly components of PSI or PSII but that are associated with photosynthesis were also detected. Transcripts in this category included *slr0228/ftsH2* encoding the FtsH2 protease crucial for the replacement of photodamaged D1 protein during the PSII repair cycle (Silva *et al*., 2003; Komenda *et al*., 2010), *ssl0020/petF* encoding the major plant-like ferredoxin Fed1 (Mazouni *et al*., 2003)*, sll0199/petE* encoding plastocyanin and *ssr2016* encoding the *Synechocystis* Pgr5 (proton gradient regulation 5) homolog. In plants and cyanobacteria, Pgr5 contributes to cyclic electron flow from PSI to the plastoquinone pool (Yeremenko *et al*., 2005; Dann and Leister, 2019), electron transfer that is associated with *crhR* expression (Kujat and Owttrim, 2000; Ritter *et al*., 2020). Another gene in this category was *trxA* (*slr0623*), encoding thioredoxin A, the most abundant of the four thioredoxins in *Synechocystis* (Hishiya *et al*., 2008), and suggests CrhR is involved in maintaining intracellular redox status by regulating protein expression associated with scavenging reactive oxygen species (ROS).

Light harvesting is an integral aspect of photosynthesis, the primary contributor to which are phycobilisomes in cyanobacteria. Among several mRNAs encoding phycobilisome (PBS) proteins in the dataset (**Table 1**) was *apcC*/*ssr3383* encoding the PBS 7.8 kDa linker polypeptide. The interacting RNA sequence encompassed 146 nt of the 5’ UTR and 54 of the 67 *apcC* codons **(Fig. 4)**, importantly ending at the start of the region from which the regulatory sRNA ApcZ originates. ApcZ targets the *ocp*/*slr1963* transcript encoding the water-soluble orange carotenoid protein OCP (Zhan *et al*., 2021). Again, interaction of this transcript with CrhR is associated with redox homeostasis since OCP functions to quench excess excitation energy absorbed by phycobilisomes, and also directly scavenges singlet oxygen ROS (^1^O_2_) (Kerfeld *et al*., 2003). Downstream of the photosynthetic electron transport chain (ETC), CrhR interaction with the *slr0148* transcript was detected (**Table 1**). The gene *slr0148* encodes the ferredoxin *fed5* (Angeleri *et al*., 2018), homologs of which are widely distributed in cyanobacteria and plants (Cassier-Chauvat and Chauvat, 2014). The Fed2 to Fed9 gene family functions to distribute electrons to a range of metabolic pathways each of which are associated with cyanobacterial tolerance to a different but overlapping range of environmental stresses (Cassier-Chauvat and Chauvat, 2014). In addition, *slr0148* is the terminal gene of 5 consecutive genes *slr0144* to *slr0148,* all of whose mRNA fragments were enriched by co-IP with CrhR_K57A_ (**Table S5**). These five genes are also jointly down-regulated after cold shift, together with the two following genes *slr0149* to *slr0151* (Georg *et al*., 2019).

**Fig. 4.**
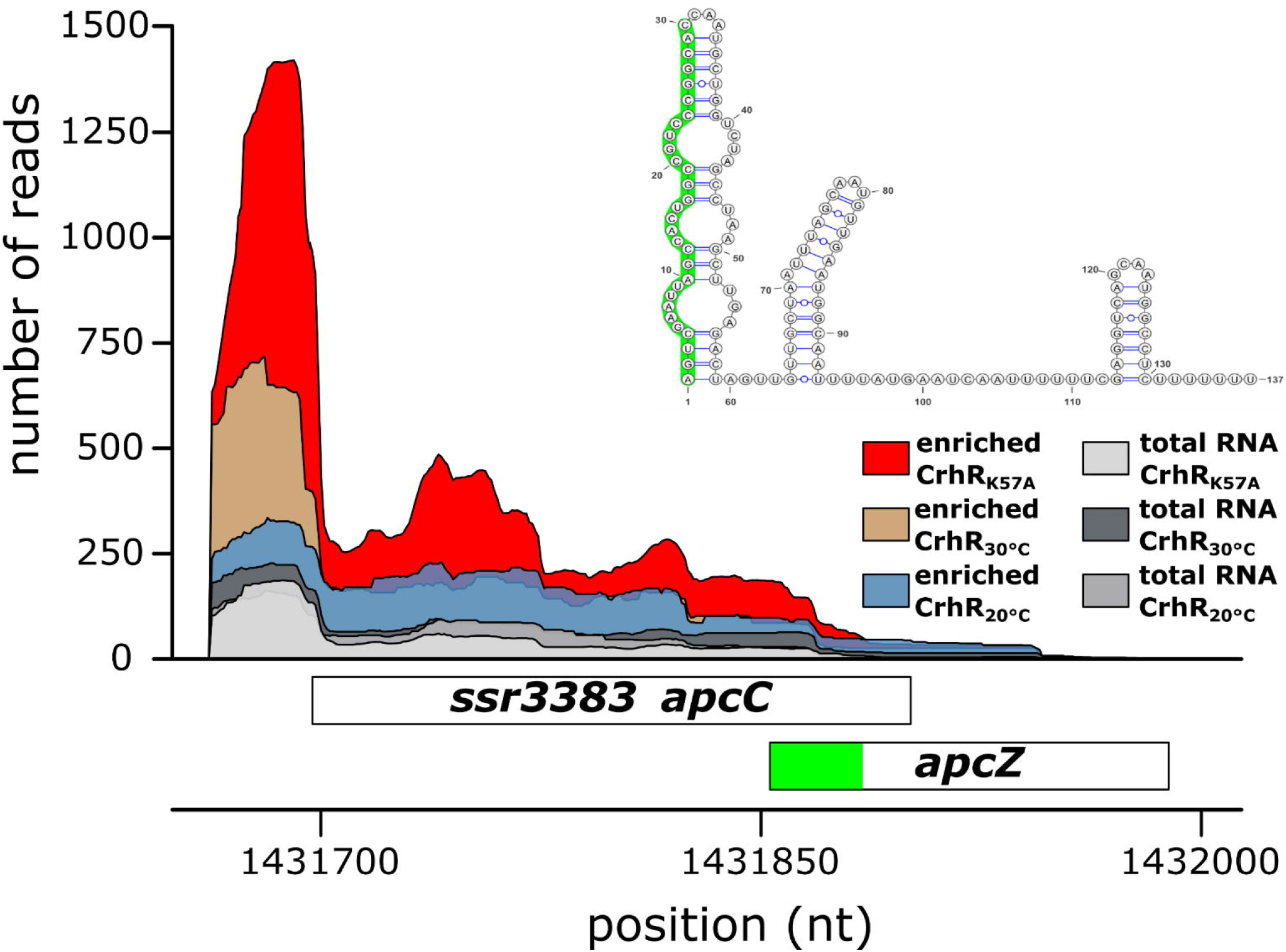
CrhR and *apcC* transcript interaction. Reads recovered from CrhR co-IP experiments cover the 5’ UTR and almost the entire coding sequence of *apcC/ssr3383* that encodes the 7.8 kDa allophycocyanin-associated linker polypeptide ApcC. Only the reads recovered in experiment 3 using CrhR_K57A_ as bait protein were significantly enriched (red) but an enrichment was also observed in CrhR_WT_ cells grown at 30°C (brown). The sRNA ApcZ originates from within the *apcC*coding sequence (boxed). The 5’ end of ApcZ was previously mapped to position 1431853, 46 nt upstream of the *apcC* stop codon (Zhan *et al*., 2021). The secondary structure of ApcZ is shown in the inset. The interaction decreases in the region from which ApcZ originates, colored light green in both the transcript and in the corresponding segment of ApcZ secondary structure where it corresponds to half of the first helix of the predicted secondary structure (nt 1 to 30). The RNAfold algorithm was used for the secondary structure prediction (Gruber *et al*., 2015).

In addition, we identified CrhR-interacting transcripts of genes more distantly related to photosynthesis, among them *slr0374* **(Fig. 3C)**, encoding the stress-responsive AAA+ protease protein Ycf46 (Singh and Sherman, 2002). Homologs of Ycf46 are highly conserved in all cyanobacterial lineages and most algal chloroplast genomes. Ycf46 proteins are involved in the regulation of CO_2_ utilization in photosynthesis but the precise function is unknown (Jiang *et al*., 2015). Transcript peaks were also recovered for *slr1674*, encoding a DUF760-containing protein described to be associated with the thermal acclimation of PSII (Rowland *et al*., 2010).

Identification of photosynthesis-associated transcripts is consistent with the published photosynthetic acclimation and loss-of-function phenotypes observed in *crhR* deletion and inactivation mutants upon temperature downshift (Rosana *et al*., 2012b; Sireesha *et al*., 2012). One should note that most of these transcripts are abundant and might be easier detected in our co-IP approach than less abundant transcripts. However, not all identified transcripts are abundant, e.g. the transcripts for genes such as *slr0442* encoding a ‘target gene of Sycrp1’ or *slr0587* belong to the set of less abundant transcriptional units according to the RNA-Seq analyses by Kopf *et al*. (2014). Nevertheless, while we detected some low abundance transcripts, the nature of the experiments and rigorous exclusion criteria employed would naturally have resulted in the exclusion of some lower expressed transcripts.

### CrhR interactome comparison with omic data

Here, we recovered in the three experiments 119 peaks belonging to 90 different genes. Previously, differential gene expression due to downshifts in temperature and the molecular effects of *crhR* deletion or inactivation have been studied at transcriptome (Prakash *et al*., 2010; Georg *et al*., 2019) and proteome level (Rowland *et al*., 2011), which generated differing regulons. Indeed 68/119 recovered peaks belong to genes for which significant differential regulation due to temperature downshifts were described at RNA level, including several transcripts encoding PSI proteins. The here identified mRNAs *psaAB, psaD, psaE* and *psaL* belong to some of the most strongly down-regulated genes at the lower temperature (see **Tables S3 to S5** for details).

Previous 2D gel electrophoresis of soluble proteins identified 16 proteins differentially expressed in the Δ*crhR* mutant at 34°C and 25 proteins at 24°C (Rowland *et al*., 2011). We found that 15 of the 90 different genes for which mRNAs were enriched in the CrhR co-IP had been identified in the analysis by Rowland *et al*. (2011) as differently expressed at protein level. This includes several proteins of the light-harvesting system including our detection of *apcA, B* and *E* transcripts coding for allophycocyanin subunits, *cpcA* and *B* encoding phycocyanin subunits and *apcC, cpcC1* and *cpcG1* encoding the linker polypeptides ApcC, CpcC1 and CpcG1 as well as the cold-inducible OCP.

Another category of proteins identified in the analysis by Rowland *et al*. (2011) are those involved in protein production and gene expression. Corresponding to the proteome described by Rowland *et al*. (2011), we identified the CrhR-interacting transcripts encoding GroEL-1 and GroEL-2 chaperones, ribosomal proteins Rps1a, Rps4, Rps14, Rpl21, Rpl28 and Slr0923 and the hypothetical protein Slr0552 (transcribed in a dicistron with *slr0551* encoding RNase J). Finally, the additional strong upregulation of the cold-inducible *slr0082* encoding methylthiotransferase RimO in Δ*crhR* was described (Rowland *et al*., 2011). While we did not detect the *slr0082* mRNA in our CrhR co-IPs, its upregulation can be explained via an operon discoordination mechanism that depends on CrhR (see below).

### Transcripts encoding proteins involved in housekeeping functions and RNA metabolism as potential CrhR targets

Another category of transcripts enriched in the CrhR co-IPs were related to house-keeping functions of the cell. Examples included *groEL1/slr2076* and *groEL2/sll0416* encoding the 60 kDa chaperonins 1 and 2 that match previous reports on the CrhR-dependent upregulation of *groEL1* and *groEL2* in *Synechocystis* (Prakash *et al*., 2010). Additionally, transcripts encoding the ribosomal proteins, *rplU/slr1678* and *rpl26/ssr1604* for the 50S ribosomal proteins L21 and L28, and *rps14/slr0628* and *rps1a/slr1356* for the 30S ribosomal proteins S14 and S1a, belonging to four separate operons, were detected. Of specific interest in relation to *crhR* expression, a transcript enriched specifically from cold stressed cultures was *desC*/*sll0541* encoding the Δ9 fatty acid desaturase DesC was identified as a potential CrhR target.

A regulatory gene of interest is *sigG/slr1545* encoding the Type 3 alternative sigma factor SigG (Huckauf *et al*., 2000), especially because it was recovered in all three experiments suggesting a stable interaction with CrhR (**Fig. 3A**). SigG is associated with the oxidative stress response caused by high light (Huckauf *et al*., 2000) and is ectopically upregulated in a Δ*rpaA* mutant in the dark (Köbler *et al*., 2018). Furthermore, interacting transcripts encoding enzymes associated with RNA turnover and metabolism were identified, including *rnj/slr0551* encoding ribonuclease J, an enzyme possessing both endonuclease and a robust 5’-exoribonuclease activity (Cavaiuolo *et al*., 2020), *slr0053* encoding the rRNA maturation RNase YbeY and, as an enzyme of RNA metabolism, *crhR* itself.

### The *crhR* mRNA is a target for auto-regulation through CrhR

The *crhR* transcript was consistently enriched in all three *crhR* lines irrespective of temperature (**Fig. 5**). The enriched peak regions within the *crhR* transcript identify three potential regions of preferred interaction: the major sequence overlaps the translational start codon and extends 216 to 270 nt into the coding region, a second region from approximately codon 125 to 219 and a third extending from approximately codon 350 to 450 of the 492 codon ORF.

**Fig. 5.**
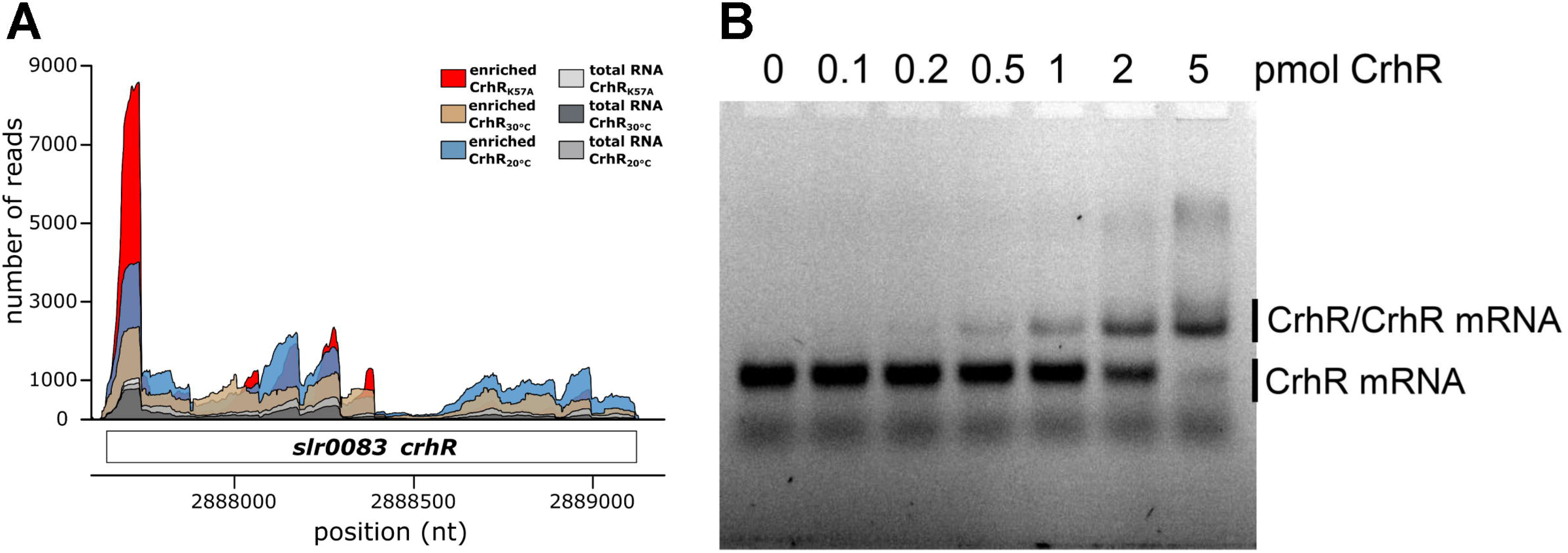
CrhR interaction with its own transcript *in vivo* and *in vitro*. **(A)** Enrichment of *crhR* mRNA in the UV-crosslinking RNA co-IP from *Synechocysti*s CrhR_WT_ grown at 20°C or 30°C (experiments 1 and 2). The colored plots show the read coverage of the enriched RNAs, as indicated. **(B)** EMSA showing binding of recombinant His-tagged CrhR to the *crhR* mRNA. For *crhR* transcript synthesis, a 150 nt DNA fragment starting from the start codon of the *crhR/slr0083* ORF was amplified with primers EMSA_CrhR-T7_Fw (carrying the T7 promoter sequence followed by two Gs) and EMSA_CrhR-T7_Rv (**Table S1**), matching to the genomic positions 2887644 to 2887793. The resulting 150 nt transcript was Cy3 labelled. Binding of 0.2 pmol of the Cy3 labelled transcripts with the indicated amounts of purified recombinant His-tagged CrhR protein was performed in the presence of poly dI-dC (1 µg).

To confirm CrhR-*crhR* mRNA interaction *in vitro,* 150 nt of the *crhR* transcript, corresponding to the primary interacting region, was tested in an electrophoretic mobility shift assay (EMSA) using His-tagged recombinant CrhR produced in *E. coli* (**Fig. 5B**). Enhanced formation of a prominent retarded protein-RNA complex was directly proportional to CrhR concentration. We conclude that the *crhR* mRNA is a preferred target of CrhR and thus an auto-regulatory effect appears likely.

## Discussion

### Involvement of CrhR in a CrhR-dependent auto-regulatory negative feedback loop

The DEAD-box RNA helicase CrhR is one of the major cold shock proteins of *Synechocystis* (Kujat and Owttrim, 2000; Prakash *et al*., 2010; Rowland *et al*., 2011; Rosana *et al*., 2012b,a; Sireesha *et al*., 2012). Consistent with observations in many other organisms that shift to lower temperature increases expression of DEAD-box RNA helicases (Owttrim, 2013), we observed previously an increased *crhR* transcript abundance after temperature downshift to 20°C, with a log_2_FC of 1.4 in the wild type and 2.6 for the 5’ portion of the gene not deleted in a partial *crhR* inactivation mutant (Georg *et al*., 2019). The higher induction in the partial *crhR* inactivation mutant pointed at a possible feedback effect of CrhR that was active in the wild type but absent in the mutant.

Regulation of *crhR* expression is complex, involving transcriptional and post-transcriptional effects (Rosana *et al*., 2012a), and an auto-regulated operon discoordination and processing mechanism was proposed (Rosana *et al*., 2020). Complicating the regulation, *crhR* is encoded in the *rimO*/*slr0082-crhR/slr0083* (*rimO-crhR*) dicistronic operon (Rosana *et al*., 2012a). However, in wild type *Synechocystis* cells, accumulation of the full-length dicistronic transcript is essentially undetectable; *rimO* and *crhR* mRNAs co-occur predominantly as monocistronic transcripts (Rosana *et al*., 2012a, 2020). Interestingly, in the absence of functional CrhR, accumulation of the entire *rimO-crhR* operon transcript is elevated compared to wild type cells (Rosana *et al*., 2020). This observation led to the assumption that CrhR may facilitate processing of the *rimO-crhR* operon transcript, although CrhR is not necessary and sufficient for the processing activity (Rosana et al., 2020). The discovery of an RNase E processing site ∼138 nt upstream of the *crhR* coding sequence then suggested that RNase E might play a role in processing of the operon. We previously showed that the *rimO-crhR* dicistronic transcript was cleaved by RNase E ∼138 nt upstream of the *crhR* start codon *in vitro* (Rosana *et al*., 2020). Interestingly, both specific RNA sequence and structure were required for RNase E substrate recognition and cleavage. Thus, we propose that CrhR may assist RNase E by interacting with the *rimO-crhR* transcript and altering its structure to the one that favors cleavage by RNase E (**Fig. 6**). In this scenario, CrhR would auto-regulate its own expression by enhancing processing of the *rimO-crhR* operon by RNase E. Moreover, CrhR could also be involved in the further processing of monocistronic *crhR* (Rosana *et al*., 2012a, 2020). In support of this hypothesis, northern and microarray analysis previously indicated that the absence of functional CrhR RNA helicase activity enhances stabilization of *crhR* transcripts at 30 and 20°C (Georg *et al*., 2019; Rosana *et al*., 2020). From the degradation constants, we concluded that at 20°C, alteration of RNA levels in the *crhR* helicase deficient mutant could be largely explained by regulation occurring at post-transcriptional level (Georg *et al*., 2019).

**Fig. 6.**
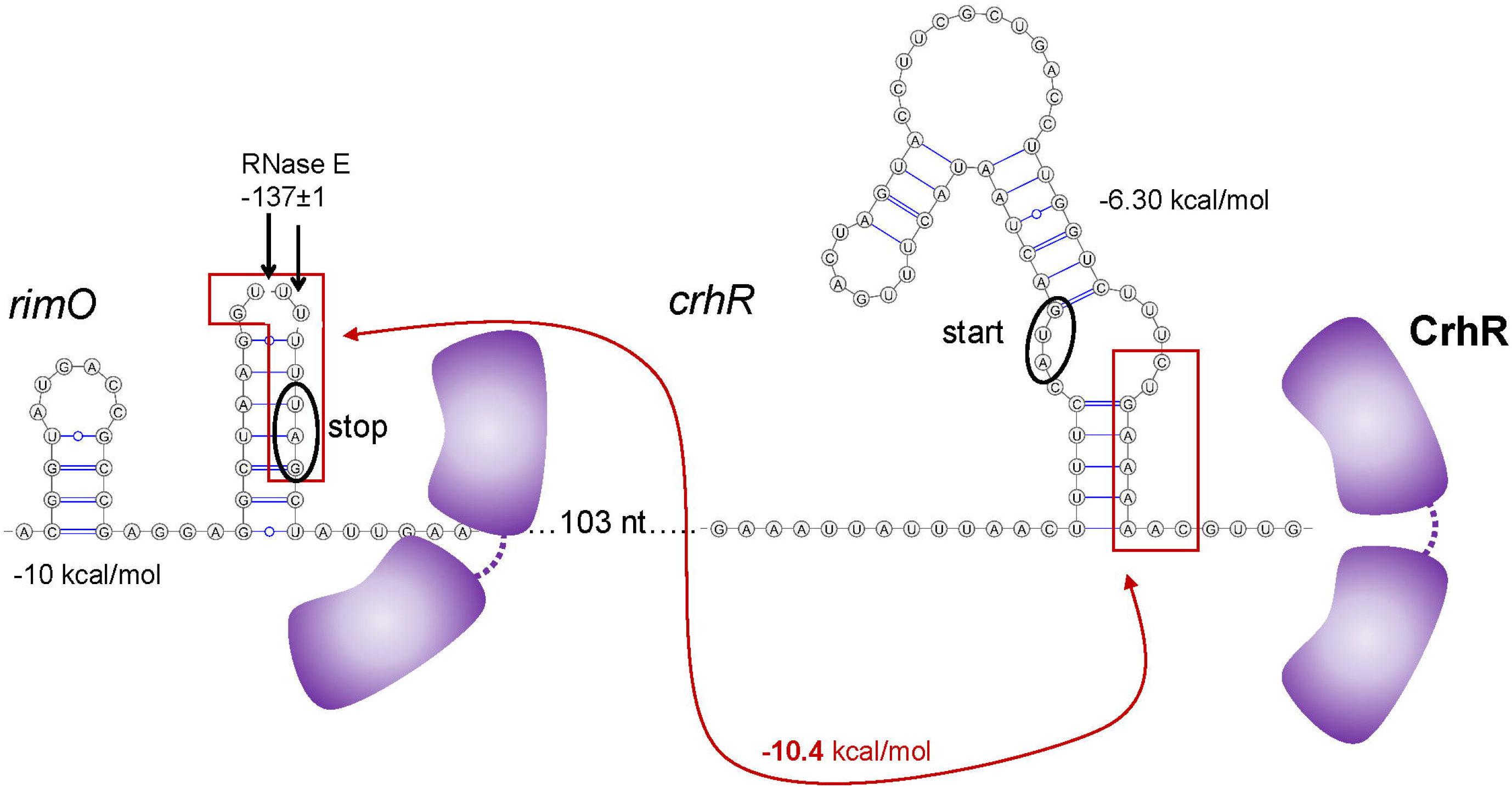
Hypothetical model for CrhR self-regulation. The translational start region of the *crhR* transcript contains a predicted secondary structure and coincides with a sharp peak of mRNA fragments recovered in all three co-IP experiments (Fig. 5A) and was verified to interact with CrhR in EMSA assays (Fig. 5B). Within this structure the AUG signal for ribosome interaction is buried (black oval) and is followed by a 9-nt sequence that is complementary to the 3’ end of the upstream *slr0082* (*rimO*) gene (red boxes). Initially, *crhR* is transcribed as a dicistronic message together with *rimO*. Operon discoordination occurs by RNase E cleavage three or four nt upstream the *rimO* stop codon within the RNase E cleavage motif (black arrows), as shown previously (Rosana *et al*., 2020). We propose that an abundance of CrhR enhances its interaction with its own mRNA by clamping the shown structure, effectively suppressing translation. Upon stress-induced activation of CrhR RNA helicase activity, this RNA secondary structure is rearranged, making the 9-nt element available for interaction with the complementary region in *rimO* (red double arrow). Further CrhR action generates the conserved single-stranded RNA motif required for processing by RNase E cleavage. The resulting monocistronic *crhR* transcript is stabilized and the detected dominating mRNA form while the remaining *rimO* transcript segment showed a drastically reduced half-life (Rosana *et al*., 2012a, 2020). Hence, CrhR is proposed to feedback regulate its own expression at two post-transcriptional levels, via operon discoordination-stabilization and at the level of translation.

In combination, these observations infer a CrhR-dependent negative feedback loop, in which binding of CrhR to *crhR* transcripts leads to auto-regulation of expression.

### The CrhR interactome *in vivo*

DEAD-box RNA helicases have been characterized as drivers and regulators of gene expression. For many DEAD-box RNA helicases, both the preferred targets as well as the mechanism of target recognition have remained unknown. This is relevant as the diversity of functions of DEAD-box RNA helicases and sequences of the auxiliary domains suggests that there is no single mechanism of specific target recognition or pathway association. Determination of the RNA helicase interactome is more challenging than for other RNA-interacting proteins since after binding, the helicase will rapidly modify the structure and dissociate. This potentially impedes the ability to co-IP interacting transcripts compared to RNA-binding proteins such as CsrA, Hfq or ProQ (Hör *et al*., 2018). To mitigate this challenge, we employed the CrhR variant, CrhR_K57A_, in which RNA residence time was predicted to be extended, since the Walker A ATP binding domain is inactivated.

This consideration was supported by the enhanced recovery of interacting transcripts from the CrhR_K57A_ strain, i.e. experiment 3 (**Fig. 1**) suggesting that RNA binding was most efficient in the CrhR_K57A_ strain. Moreover, the partial complementation of the Δ*crhR* phenotype by CrhR_K57A_ (**Fig. 2**) supported the view that this variant had reduced RNA helicase enzymatic activity. The overlap in the enriched targets identified in the three experiments consists of transcripts belonging to only four genes (*crhR, psaA, sigG* and *ycf46*) as shown in the Venn diagram in **Fig. S2**.

Functionally, the prime targets of CrhR interaction were consistently observed to include transcripts associated with photosynthesis, light harvesting or directly encoding core photosystem proteins (**Table 1**). These findings are consistent with observations that the rapid cessation of photosynthesis upon temperature downshift from 30 to 20°C is the major physiological consequence created by the absence of CrhR RNA helicase activity (Rosana *et al*., 2012b). A vital aspect relevant to photosynthesis is the reported inability of the Δ*crhR* mutant to properly perform light-induced state transitions, which occur as part of the normal acclimation response to downshifts in temperature in the wild type (Sireesha *et al*., 2012). Our data provide deeper insight into the molecular basis of these observations. In contrast to the monomeric PSI structures in plants and algae, PSI in many cyanobacteria is mainly trimeric, in some strains tetrameric (Li *et al*., 2019). The trimeric structure enlarges the antenna system under low-light conditions, while the dynamic regulation of the PSI trimer-to-monomer ratio impacts the speed of state transitions (Aspinwall *et al*., 2004). The PSI reaction center protein subunit XI encoded by *psaL* is crucial for the multimerization status of PSI (Aspinwall *et al*., 2004; Li *et al*., 2019). Here, we observed a pronounced interaction between CrhR and a segment of the *psaL* mRNA that contains the 5’ UTR and the first 30 codons (**Fig. S3**). This region furthermore contains the interaction site for the photosynthesis regulatory RNA1, PsrR1 (Georg *et al*., 2014). Previous analyses demonstrated that the proportion of PSI trimers is reduced in Δ*crhR* relative to wild type upon temperature downshift (Sireesha *et al*., 2012). Therefore, the interaction between the cold-induced CrhR and this region is predicted to be necessary for the required expression of *psaL,* likely by facilitating translation, as diagramed in **Fig. S3**.

A role of CrhR in the biology of photosynthesis-relevant mRNAs is further consistent with the demonstrated localization of ribosome-associated CrhR at the thylakoid membrane (Rosana *et al*., 2016) and the observation that *psbA* and *psaA* mRNAs are transported to the thylakoid membrane and possibly translated by thylakoid-associated ribosomes (Tyystjärvi *et al*., 2001; Mahbub *et al*., 2020). These findings point to a possible involvement of CrhR in physiological secondary structure rearrangements of the identified mRNAs that could influence translation on thylakoid membrane associated ribosomes.

Since *Synechocystis* encodes only a single DEAD-box RNA helicase, it is likely that CrhR operates in multiple pathways. These roles could be associated with different stresses, independent of temperature, as observed for the *Staphylococcus aureus* helicase, CsdA (Oun *et al*., 2013; Khemici *et al*., 2020). Transcripts recovered for ribosomal proteins and ribonucleases support this idea. Indeed, a role in translation regulation is supported by the observation that CrhR interacts with the *rps1a/slr1356* transcript encoding Rps1a. In previous analyses, Rps1a sedimented away from the majority of ribosomal proteins (Riediger *et al*., 2021) and was reported to perform a role in the Shine-Dalgarno-independent initiation of translation (Mutsuda and Sugiura, 2006; Nakagawa *et al*., 2010), another process which may hence be related to CrhR helicase activity.

Another intriguing example for the possible involvement of CrhR in functionally relevant secondary structure re-arrangements was observed for *apcC/ssr3383* where the recovered reads terminated in the region from which the regulatory sRNA ApcZ originates. ApcZ is an sRNA, which targets the *ocp/slr1963* mRNA encoding the water-soluble OCP by inhibiting its translation (Zhan *et al*., 2021). If expressed, OCP directly senses light intensity and induces thermal energy dissipation under stress conditions (Muzzopappa and Kirilovsky, 2020). Therefore, its expression is tightly controlled and, in this context, ApcZ can inhibit or delay the production of OCP. This regulation is also relevant under nitrogen starvation conditions, when ApcZ is induced from a transcription start site (TSS) that is activated by the transcription factor NtcA (Zhan *et al*., 2021). However, ApcZ is also detectable as a transcript under other conditions where it derives from processing of the *apcABC* mRNA. Interestingly, the co-IP coverage included the first 30 nt of ApcZ, matching precisely the upstream portion of the helical structure located at the 5’ end of ApcZ (**Fig. 4**). Therefore, it is tempting to speculate that CrhR interaction and helicase activity is either required for i) ApcZ processing, ii) to open the ApcZ structure for interaction, or possibly iii) to facilitate ApcZ-*ocp* duplex formation. These functions are supported by the observation that the *ocp* mRNA was identified as a potential CrhR target in our co-IP experiment (**Table 1**), that it accumulates at a higher level upon temperature downshift (Georg *et al*., 2019) and by the ability of CrhR to catalyze both RNA duplex unwinding and annealing (Chamot *et al*., 2005). In conjunction, the OCP protein level was significantly increased at the lower temperature (Rowland *et al*., 2011).

### Overlaps in the regulon controlled by RpaB and the set of transcripts interacting with CrhR

Several of the genes whose transcripts interacted with CrhR were previously predicted or demonstrated to play a role in redox signaling or are regulated in a redox-dependent manner. This applies first and foremost to *crhR* itself (Kujat and Owttrim, 2000; Ritter *et al*., 2020). Another is thioredoxin A, the homolog of plant m-type thioredoxin and therefore sometimes also called TrxM, which plays a major role in redox regulation in *Synechocystis* (Hishiya *et al*., 2008). A protein interacting with TrxM *in vitro* is the redox-responsive transcription factor RpaB (Kadowaki *et al*., 2015). RpaB recognizes the “high light regulatory 1’’ (HLR1) promoter element, a pair of imperfect 8-nt long direct repeats (G/T)TTACA(T/A) (T/A) separated by two random nucleotides (Eriksson *et al*., 2000; Kappell and van Waasbergen, 2007). Binding of RpaB to HLR1 promoter motifs yields one of two different outcomes, repression or activation under low light depending on the distance between the location of these elements and the TSS (reviewed by Riediger *et al*., 2018). RpaB is the response regulator (RR) of the histidine kinase Hik33 in a two-component signal transduction system that controls >150 target promoters in *Synechocystis*, regulating expression of diverse genes associated with photosynthesis (Riediger *et al*., 2019). Thus, there is a striking correspondence that approximately two thirds of the transcripts recovered in our CrhR-co-IP analysis belong to the RpaB regulon (**Table 1**). Identification of RpaB regulon members as CrhR-interacting transcripts therefore suggests a scenario in which CrhR contributes to the post-transcriptional regulation of RpaB regulon expression. Together, these observations support the function of CrhR as an important post-transcriptional regulatory element within the network of redox-dependent signaling and regulation.

## Acknowledgements

We thank the Freiburg Galaxy Team for maintaining this great bioinformatics resource.

## Author contributions

WRH designed the study, CS established the cDNA library preparation and supported the bioinformatic analysis; JF, BH and RR sequenced cDNA libraries; AM, FH, RB and WRH analyzed data; AS and JSSP constructed the *Synechocystis* strains. All other experiments were carried out by AM. AM, GWO and WRH wrote the paper with contributions from all authors.

## Data availability

The RNA-seq data and raw sequence data from co-IP analyses have been deposited in the SRA database https://www.ncbi.nlm.nih.gov/sra/ and are openly available under the accession numbers SAMN14615974 to SAMN14615989 and SAMN14651279 to SAMN14651286.

