## Supplemental data for "The temperature-regulated DEAD-box RNA helicase CrhR interactome: Autoregulation and photosynthesis-related transcripts"

### Supplementary Material

### Supplementary Tables

**Table S1. Synthetic DNA and RNA oligonucleotide primers used in this work.** *Restriction sites are underlined and indicated by a vertical arrow. The T7 RNA polymerase promoter sequence is in boldface letters and underlined. To barcode the different libraries, PCR primer 2 variants were used with different hexanucleotides at the indicated N6 position.

| **Primer name** | **Sequence (5' to 3' direction)** |
|---|---|
| 3xFL-CrhR-F1 | GATTACAAGGATGACGATGACAAGACTAATACTTTGACTAGTACCTTCG |
| 3xFL-CrhR-F2 | GATTATAAAGATCATGATATCGATTACAAGGATGACGATGACAAGAC |
| 3xFL-CrhR-F3 | GCCA↓TATGGACTACAAAGACCATGACGGTGATTATAAAGATCATGATATC (*Nde*I*) |
| 3xFL-CrhR-R | CCGT↓CTAGACCAGAGCT↓CATTACTGTTGGCGATCACTATAGGCAGGACG (*Xba*I, *Sac*I*) |
| petE-FP | TTTCAGCAAGATCGG CAAGGCAAAACCACCGTTATCAGCAG |
| petE-RP | TTTGTAGTccATATGACTTCTTGG CGATTGTATCTATAGGGACTTG |
| rrnB-TT-F | GGCGAGCT↓CGCCTGGCGGCAGTAGCGCGG (*Sac*I*) |
| rrnB-TT-R | CCT↓CTAGAGGAATTCAAAAAGGCCATCCGTCAGGATG (*Xba*I*) |
| crhR(K57A)_inverse_fw | AATGCGGTCCATCAAAGGTAGGGCAAAGGCGGCGGTTGCCCCCGTGCCGGTTTGGG |
| crhR(K57A)_inverse_rv | GACCCAGAGGGGGACTTGCAAGCCTTAATTTTGACC |
| petE-3xF-rrnb_neg_ctrl_FP | GCCGAGCT↓CGCCTGGCGGCAG (*Sac*I*) |
| petE-3xF-rrnb_neg_ctrl_RP | GGCGAGCT↓CTTACTTGTCATCGTCATCCTTGTAATCGATATCA (*Sac*I*) |
| petE_foR-pVZ_FP | CGGG↓AATTCCAAGGCAAAACCACCGTTATCAGCAG (*Eco*RI*) |
| rrnB_foR-pVZ_RP | CGGG↓AATTCAAAAAGGCCATCCGTCAGGATG (*Eco*RI*) |
| pQE70_crhR-fw | GAAATTAAGC ATGACTAATACTTTGACTAGTACC |
| pQE70_crhR-rv | TGGTGATGAGATCT CTGTTGGCGATCACTATAGG |
| pQE_GIBSOn-fw | AGATCTCATCACCATCACCATC |
| pQE_Gibson-rv | GCTTAATTTCTCCTCTTTAATGAATTCTGTG |
| EMSA_CrhR-T7_Fw | **TAATACGACTCACTATAGG**ATGACTAATACTTTGACTAGTACCTTC |
| EMSA_CrhR-T7_Rv | TTGGGCCAGCATGTCCCG |
| Loop primer | AGATCGGAAGAGAGACGTGTGCTCTTCCGATCTNNNNNNN (N = A,C,G, or T) |
| UGA linker | CCCUACACGACGCUCUUCCGAUCUGA |
| PCR primer 1 | AATGATACGGCGACCACCGAGATCTACACTCTTTCCCTACACGACGCTCTTCCGATCTGA |
| PCR primer 2 | CAAGCAGAAGACGGCATACGAGAT[N6]GTGACTGGAGTTCAGACGTGTGCTCTTCCGATCT |

**Table S2.** Read numbers from the *in vivo* UV-crosslinking of RNA to CrhR and to $CrhR_{K57A}$ in the three performed experiments (cf. **Fig. 1**). Absolute and relative numbers of reads derived from RNA-seq data analysis.

| Sample | Total number | Mapped to chromosome | % of mapped |
|---|---|---|---|
| **20°C** | | | |
| **Experiment 1: UV-crosslinking RNA pulldown from *Synechocystis* Δ*crhR*/FLAG-CrhR and controls** | | | |
| **FLAG-CrhR 1** | 18,387,764 | 17,126,280 | 93.1 |
| **FLAG-CrhR 2** | 16,744,574 | 15,900,654 | 95.0 |
| **Δ*crhR*/FLAG-CrhR Total RNA 1** | 20,564,614 | 19,399,684 | 94.3 |
| **Δ*crhR*/FLAG-CrhR Total RNA 2** | 23,425,924 | 22,037,360 | 94.1 |
| **FLAG 1** | 21,778,800 | 20,312,372 | 93.3 |
| **FLAG 2** | 20,769,842 | 19,371,524 | 93.3 |
| **Δ*crhR*/FLAG Total RNA 1** | 19,238,410 | 17,852,380 | 92.8 |
| **Δ*crhR*/FLAG Total RNA 2** | 17,592,288 | 16,194,922 | 92.1 |
| **30°C** | | | |
| **Experiment 2: UV-crosslinking RNA pulldown from *Synechocystis* Δ*crhR*/FLAG-CrhR and controls** | | | |
| **FLAG-CrhR 1** | 15,817,350 | 14,457,262 | 91.4 |
| **FLAG-CrhR 2** | 19,740,942 | 18,694,828 | 94.7 |
| **Δ*crhR*/FLAG-CrhR Total RNA 1** | 23,761,732 | 22,485,096 | 94.6 |
| **Δ*crhR*/FLAG-CrhR Total RNA 2** | 20,837,756 | 19,667,718 | 94.4 |
| **FLAG 1** | 20,373,452 | 19,151,100 | 94.0 |
| **FLAG 2** | 18,615,012 | 17,386,892 | 93.4 |
| **Δ*crhR*/FLAG Total RNA 1** | 14,340,512 | 13,387,226 | 93.4 |
| **Δ*crhR*/FLAG Total RNA 2** | 21,776,284 | 20,140,512 | 92.5 |
| **Experiment 3: UV-crosslinking RNA pulldown from *Synechocystis* $CrhR_{K57A}$ and controls** | | | |
| **FLAG-CrhR(K57A) 1** | 15,603,510 | 14,457,362 | 92.7 |
| **FLAG-CrhR(K57A) 2** | 28,388,296 | 26,368,740 | 92.9 |
| **Δ*crhR*/FLAG-CrhR(K57A) Total RNA 1** | 30,248,016 | 28,194,156 | 93.2 |
| **Δ*crhR*/FLAG-CrhR(K57A) Total RNA 2** | 29,340,520 | 27,046,542 | 92.2 |
| **FLAG 1** | 12,416,532 | 11,556,000 | 93.1 |
| **FLAG 2** | 28,490,174 | 26,462,782 | 92.9 |
| **Δ*crhR*/FLAG Total RNA 1** | 43,713,322 | 40,553,504 | 92.8 |
| **Δ*crhR*/FLAG Total RNA 2** | 33,298,496 | 30,744,948 | 92.3 |

**Table S3. Transcripts enriched in the UV-crosslinking RNA pulldown from *Synechocystis* CrhR$_{WT}$ grown at 20°C (Experiment 1 in Fig. 1).** The identified peaks are numbered (P), their start and end is given in nt according to Genbank file NC_000911 followed by information indicating if the respective transcripts originated from the forward (+) or reverse strand (-). The enrichment is given as $log_2$FC together with the adjusted p value ($p_{adj}$). The IDs of the respective transcript follow the nomenclature in Genbank file NC_000911 and indicate if the peak was within the coding region (CDS) or included also part of an UTR. The final two rows give the annotation of the CrhR-interacting RNAs and information on their regulation in previous transcriptome (1) (Georg *et al.*, 2019) or proteome (2) (Rowland *et al.*, 2011) studies of *crhR* mutants. The experiment was performed in biological duplicates.

| P | Start | End | S | $log_2$FC | $p_{adj}$ | ID | Annotation | Regulation |
|---|---|---|---|---|---|---|---|---|
| 1 | 890731 | 890985 | + | 1.23 | 1.5E-07 | ssr2016 CDS | Pgr5, soluble electron acceptor | strongly cold-induced at RNA level (1) |
| 2 | 2529108 | 2529338 | + | 1.18 | 8.1E-09 | slr0228 CDS | FtsH2 protease | lower RNA level in Δ*crhR* at both temperatures (1) |
| 3 | 1167439 | 1167768 | + | 1.06 | 3.9E-03 | smr0009 CDS | PSII reaction center protein PsbN | - |
| 4 | 2888690 | 2889127 | + | 1.05 | 3.7E-09 | slr0083 CDS, 3'UTR | DEAD-box RNA helicase CrhR | cold induction of *slr0082*, the first gene in this operon, was observed at protein level in Δ*crhR* (2) |
| 5 | 318624 | 318767 | + | 0.97 | 4.0E-05 | ssr1604 CDS, 5'UTR | ribosomal protein L28 | consistent with reported effects on ribosomal proteins (2) |
| 6 | 983689 | 984074 | + | 0.96 | 2.0E-07 | slr1545 CDS, 5'UTR | extracytoplasmic function (ECF) sigma factor SigG | lower RNA level in the cold in WT and in Δ*crhR* (1) |
| 7 | 1954655 | 1954908 | + | 0.95 | 9.2E-04 | slr1674 CDS, 5'UTR, | Thermoprotector protein of PSII | - |
| 8 | 2576009 | 2576295 | + | 0.91 | 3.0E-05 | ssr0109 CDS, 5'UTR; slr0053 CDS | uncharacterized 62 aa protein; putative rRNA maturation RNase YbeY | - |
| 9 | 2887625 | 2887994 | + | 0.89 | 6.9E-11 | slr0083 CDS | DEAD-box RNA helicase CrhR | cold induction of *slr0082*, the first gene in this operon, was observed at protein level in Δ*crhR* (2) |
| 10 | 2219257 | 2219424 | + | 0.85 | 4.8E-03 | slr1926 CDS, 5'UTR | hypothetical protein | RNA level is cold-induced in WT and in Δ*crhR* (1) |
| 11 | 1096987 | 1097148 | + | 0.81 | 8.7E-04 | slr1356 CDS, 5'UTR | ribosomal protein S1a | cold induction at protein level in Δ*crhR* (2) and consistent with reported effects on ribosomal proteins (2) |

| | | | | | | | | |
|---|---|---|---|---|---|---|---|---|
| 12 | 2961098 | 2961423 | + | 0.76 | 1.3E-02 | slr0623 CDS, 5'UTR | thioredoxin A | RNA level is cold-repressed in WT and in Δ*crhR* (1) |
| 13 | 2098723 | 2098941 | + | 0.75 | 1.2E-03 | ncr1010, slr0447 CDS, 5'UTR | Ncr1010; periplasmic protein, ABC-type urea transport system substrate-binding protein UrtA | RNA level higher in Δ*crhR* than in WT, but cold-repressed in both strains (1) |
| 14 | 3457402 | 3457655 | + | 0.73 | 1.8E-02 | slr1655 5'UTR, CDS | PSI subunit XI, PsaL | strong down-regulation at RNA level at the lower temperature (1); consistent with lower PSI trimer-to-monomer ratio in the cold |
| 15 | 2484892 | 2485282 | - | 0.73 | 8.1E-09 | ssl0020 5'UTR, CDS | ferredoxin I, PetF | slight down-regulation in the cold at RNA level (1) |
| 16 | 3253100 | 3253502 | + | 0.70 | 9.3E-03 | slr0552 CDS | hypothetical protein | divergent regulation in Δ*crhR* versus WT reported at protein level (2) |
| 17 | 2798031 | 2798246 | + | 0.65 | 2.7E-02 | slr0923 CDS, 5'UTR, | YCF65 | Induction in the cold at RNA (1) and protein level (2), both more pronounced in WT; possibly ribosome-associated |
| 18 | 2888016 | 2888299 | + | 0.65 | 5.1E-05 | slr0083 CDS | DEAD-box RNA helicase CrhR | cold induction of *slr0082*, the first gene in this operon, was observed at protein level in Δ*crhR* (2) |
| 19 | 303507 | 303756 | + | 0.63 | 3.6E-15 | ssr2153 CDS | uncharacterized 65 aa protein, induced at low $CO_2$ | - |
| 20 | 2050396 | 2050657 | + | 0.63 | 4.0E-02 | slr1638 CDS, 5'UTR | DUF760 protein | - |
| 21 | 2375994 | 2376375 | + | 0.61 | 4.6E-03 | slr0374 CDS | Ycf46 | down-regulated at RNA level at cold temperature, more pronounced in WT than in Δ*crhR* (1) |
| 22 | 1767669 | 1767988 | + | 0.61 | 4.0E-02 | slr1963 CDS | water-soluble carotenoid protein OCP | enhanced expression in the cold at RNA and protein level in both strains (1, 2) |
| 23 | 3252507 | 3252917 | + | 0.60 | 4.0E-05 | slr0551 CDS | ribonuclease J | divergent regulation of *slr0552*, second gene in this operon reported in Δ*crhR* versus WT at protein level (2) |
| 24 | 2603906 | 2604110 | + | 0.57 | 2.5E-02 | slr0469 CDS | ribosomal protein S4 | consistent with reported effects on ribosomal proteins (2); cold-induced at RNA level, more pronounced in WT (1) |

| | | | | | | | | |
|---|---|---|---|---|---|---|---|---|
| 25 | 2823761 | 2824049 | - | 0.56 | 1.3E-09 | sll0541-5'UTR | desC/des9, Acyl-CoA desaturase | - |
| 26 | 2495231 | 2495441 | - | 0.51 | 4.0E-02 | sll0006 CDS, 5'UTR | putative aminotransferase AspC | - |
| 27 | 959370 | 959752 | + | 0.49 | 4.9E-02 | slr1841 CDS | porin, major outer membrane protein | - |
| 28 | 2791426 | 2791783 | + | 0.28 | 3.4E-11 | slr0915 CDS, 3'UTR; 6803t34-exon 2; slr0917 5'UTR | tRNA(Leu)$^{UAA}$ exon 2, endonuclease; *bioF*, 8-amino-7-oxononanoate synthase | - |

**Table S4. Transcripts enriched in the UV-crosslinking RNA pulldown from *Synechocystis* CrhR$_{WT}$ grown at 30°C (Experiment 2).** Column titles and acronyms as in **Table S3.**

| P | Start | End | S | log$_2$FC | $p_{adj}$ | ID | Annotation | Regulation |
|---|---|---|---|---|---|---|---|---|
| 1 | 1015575 | 1015843 | + | 1.94 | 4.8E-02 | slr1751 CDS | CtpC, periplasmic carboxyl-terminal protease | downregulated in the cold at RNA level (1) |
| 2 | 2888696 | 2888995 | + | 1.80 | 1.6E-03 | slr0083 CDS | DEAD-box RNA helicase CrhR | cold induction of *slr0082*, the first gene in this operon, was observed at protein level in Δ*crhR* (2) |
| 3 | 2529057 | 2529411 | + | 1.79 | 1.6E-02 | slr0228 CDS | FtsH2 protease | somewhat upregulated in the cold in both strains at RNA level (1) |
| 4 | 3251635 | 3251903 | + | 1.69 | 3.7E-02 | slr0551 CDS | ribonuclease J | divergent regulation of *slr0552*, second gene in this operon reported in Δ*crhR* versus WT at protein level (2) |
| 5 | 2575990 | 2576292 | + | 1.69 | 1.3E-02 | ssr0109 5'UTR, slr0053 CDS | uncharachterized 62 aa protein; putative rRNA maturation RNase YbeY | - |
| 6 | 3252662 | 3252914 | + | 1.59 | 4.8E-02 | slr0551 CDS | ribonuclease J | divergent regulation of *slr0552*, second gene in this operon reported in Δ*crhR* versus WT at protein level (2) |
| 7 | 2376063 | 2376367 | + | 1.57 | 8.4E-06 | slr0374 CDS | Ycf46 | down-regulated at RNA level at cold temperature, more pronounced in WT than in Δ*crhR* (1) |
| 8 | 983777 | 984044 | + | 1.42 | 4.6E-04 | slr1545 5'UTR, CDS | extracytoplasmic function sigma factor SigG | lower RNA level in the cold in WT and in Δ*crhR* (1) |
| 9 | 2888062 | 2888418 | + | 1.30 | 7.9E-03 | slr0083 CDS | ORF of crhR, DEAD-box RNA helicase CrhR | cold induction of *slr0082*, the first gene in this operon, was observed at protein level in Δ*crhR* (2) |
| 10 | 2887625 | 2887895 | + | 1.08 | 3.7E-02 | slr0083 5'UTR, CDS | ORF of crhR, DEAD-box RNA helicase CrhR | |
| 11 | 2526183 | 2526380 | - | 1.03 | 2.1E-04 | sll0199 CDS; ncl1260, ncl1270 | plastocyanin PetE; Ncl1260; Ncl1270 | - |
| 12 | 941580 | 941917 | + | 1.02 | 3.2E-04 | slr1834 5'UTR, CDS | PsaA, P700 apoprotein subunit Ia | lower RNA level in the cold in WT and in Δ*crhR* (1) |
| 13 | 2644753 | 2645020 | + | 0.99 | 3.1E-03 | slr0628 5'UTR, CDS | 30S ribosomal protein S14 | consistent with reported effects on ribosomal proteins (2); cold |

| | | | | | | | | |
|---|---|---|---|---|---|---|---|---|
| | | | | | | | | induction seen in WT is almost missing in Δ*crhR* (1) |
| 14 | 303516 | 303746 | + | 0.96 | 4.4E-03 | ssr2153 CDS | 65 aa protein induced at low $CO_2$ | - |
| 15 | 942173 | 942663 | + | 0.93 | 1.4E-03 | slr1834 CDS | PSI P700 apoprotein subunit Ia PsaA | lower RNA level in the cold in WT and in Δ*crhR* (1) |
| 16 | 1329131 | 1329252 | - | 0.82 | 2.5E-02 | 6803t13-3'UTR, tRNA | tRNA(Asp)$^{GUC}$ | - |

**Table S5. RNA enriched in the UV-crosslinking RNA pulldown from *Synechocystis* CrhR$_{K57A}$ grown at 30°C (experiment 3).** Column titles and acronyms as in **Table S3.**

| P | Start | End | S | $log_2FC$ | $p_{adj}$ | ID | Annotation | Regulation |
|---|---|---|---|---|---|---|---|---|
| 1 | 1911446 | 1911828 | + | 2.95 | 2.82E-27 | slr1152 CDS | hypothetical gene | lower RNA level in the cold in WT and in Δ*crhR* (1) |
| 2 | 2427868 | 2428211 | + | 2.78 | 2.05E-24 | slr0342 5'UTR, gene | cytochrome b6 PetB | somewhat lower RNA level in the cold in WT and in Δ*crhR* (1) |
| 3 | 943649 | 944126 | + | 2.75 | 4.70E-41 | slr1834 CDS | PSI P700 apoprotein subunit Ia PsaA | lower RNA level in the cold in WT and in Δ*crhR* (1) |
| 4 | 916100 | 916489 | + | 2.73 | 8.63E-26 | slr2076 CDS | groEL1, 60 kDa chaperonin 1 | cold induction of this gene and the preceding *groES* at protein level in WT and Δ*crhR* (2) |
| 5 | 2888068 | 2888428 | + | 2.72 | 4.34E-32 | slr0083 CDS | DEAD-box RNA helicase CrhR | cold induction of *slr0082*, the first gene in this operon, was observed at protein level in Δ*crhR* (2) |
| 6 | 942606 | 943172 | + | 2.68 | 8.43E-36 | slr1834 CDS | PSI P700 apoprotein subunit Ia PsaA | lower RNA level in the cold in WT and in Δ*crhR* (1) |
| 7 | 942171 | 942587 | + | 2.67 | 1.60E-36 | slr1834 CDS | PSI P700 apoprotein subunit Ia PsaA | |
| 8 | 2178410 | 2178786 | + | 2.66 | 1.28E-34 | slr0146 CDS | PSII assembly proteins operon | |
| 9 | 2607661 | 2607952 | + | 2.64 | 1.02E-22 | ncr1350-ncRNA; slr0473 CDS | Ncr1350; phytochrome-like protein Cph1 | RNA level higher in Δ*crhR* than in WT (1), but level decreased over cold shift in both strains (1) |
| 10 | 2203869 | 2204206 | + | 2.61 | 8.00E-23 | slr0161 CDS; slr0162 CDS | twitching motility protein PilT, pilin biogenesis protein PilC | lower RNA level in the cold in WT and in Δ*crhR* (1) |
| 11 | 2177694 | 2178089 | + | 2.59 | 1.19E-24 | slr0144 CDS; slr0145 CDS | PSII assembly proteins operon | |

| | | | | | | | | |
|---|---|---|---|---|---|---|---|---|
| 12 | 2179323 | 2179758 | + | 2.58 | 5.95E-25 | slr0148- slr0147 5'UTR, CDS | PSII assembly proteins operon | |
| 13 | 916518 | 916812 | + | 2.53 | 2.25E-24 | slr2076 CDS | groEL1, 60 kDa chaperonin 1 | cold induction of this gene and the preceding *groES* at protein level in WT and Δ*crhR* (2) |
| 14 | 941540 | 941915 | + | 2.52 | 3.80E-30 | slr1834 5'UTR, CDS | PSI P700 apoprotein subunit Ia PsaA | lower RNA level in the cold in WT and in Δ*crhR* (1) |
| 15 | 2292158 | 2292507 | + | 2.39 | 2.71E-21 | slr0335-5'UTR, CDS | phycobiliprotein ApcE | |
| 16 | 2887625 | 2887825 | + | 2.32 | 2.45E-26 | slr0083 5'UTR, CDS | DEAD-box RNA helicase CrhR | cold induction of *slr0082*, the first gene in this operon, was observed at protein level in Δ*crhR* (2) |
| 17 | 946045 | 946425 | + | 2.29 | 3.99E-25 | slr1835 CDS | PSI P700 chlorophyll a apoprotein A2 PsaB | lower RNA level in the cold in WT and in Δ*crhR* (1), consistent with location as second gene in a dicistron with *psaA* |
| 18 | 2178094 | 2178408 | + | 2.28 | 9.06E-18 | slr0145 CDS | PSII assembly proteins operon | lower RNA level in the cold in WT and in Δ*crhR* (1) |
| 19 | 1431015 | 1431543 | + | 2.25 | 2.36E-23 | slr1986 CDS | allophycocyanin beta chain ApcB | |
| 20 | 2176903 | 2177136 | + | 2.22 | 1.22E-15 | slr0144 5'UTR, CDS | PSII assembly proteins operon | |
| 21 | 2479838 | 2480186 | + | 2.14 | 3.40E-22 | slr0011 CDS | Rubisco chaperonin RbcX | - |
| 22 | 959407 | 959752 | + | 2.12 | 8.70E-18 | slr1841 CDS | major outer membrane porin | - |
| 23 | 3535986 | 3536209 | + | 2.12 | 1.52E-15 | slr0587 CDS | hypothetical protein | somewhat lower RNA level in the cold in WT and in Δ*crhR* (1) |
| 24 | 916823 | 917223 | + | 2.12 | 2.49E-15 | slr2076 CDS | groEL1, 60 kDa chaperonin 1 | cold induction of this gene and the preceding *groES* at protein level in WT and Δ*crhR* (2) |

| | | | | | | | | |
|---|---|---|---|---|---|---|---|---|
| 25 | 1431552 | 1431861 | + | 2.11 | 6.79E-17 | ssr3383 CDS | PBS 7.8 kDa linker polypeptide ApcC | strongly reduced RNA levels in WT and mutant in the cold (1) |
| 26 | 1957812 | 1958225 | + | 2.11 | 1.46E-16 | slr1678 CDS | 50S ribosomal protein L21 RplU | consistent with reported effects on ribosomal proteins (2); cold induction seen in WT is almost missing in Δ*crhR* (1) |
| 27 | 2780114 | 2780469 | + | 2.08 | 2.08E-16 | slr0906 CDS | PSII CP47 reaction center protein PsbB | lower RNA levels in WT and mutant in the cold (1) |
| 28 | 2478911 | 2479284 | + | 2.05 | 1.85E-12 | slr0009 CDS | ribulose bisphosphate carboxylase large subunit RbcL | - |
| 29 | 3230578 | 3230866 | + | 1.99 | 3.35E-15 | slr0927 CDS, 3'UTR | PSII D2 protein PsbD | - |
| 30 | 1430662 | 1430991 | + | 1.94 | 1.47E-16 | slr2067 CDS | allophycocyanin alpha chain ApcA | strongly reduced RNA levels in WT and mutant in the cold (1) |
| 31 | 2480305 | 2480689 | + | 1.91 | 9.75E-18 | slr0011 CDS; slr0012 CDS | Rubisco chaperonin RbcX; ribulose bisphosphate carboxylase small subunit RbcS | - |
| 32 | 944815 | 945321 | + | 1.90 | 6.86E-15 | slr1835 CDS | PSI P700 chlorophyll a apoprotein A2 PsaB | lower RNA level in the cold in WT and in Δ*crhR* (1), consistent with location as second gene in a dicistron with *psaA* |
| 33 | 2375834 | 2376372 | + | 1.88 | 9.30E-18 | slr0374 CDS | Ycf46 | down-regulated at RNA level at cold temperature, more pronounced in WT than in Δ*crhR* (1) |
| 34 | 3230117 | 3230247 | + | 1.86 | 8.19E-12 | slr0927 CDS | PSII D2 protein PsbD | - |
| 35 | 126819 | 127109 | + | 1.84 | 6.32E-17 | slr0737 CDS, 3'UTR | PSI subunit II PsaD | strongly reduced RNA levels in WT and mutant in the cold (1) |
| 36 | 1430313 | 1430660 | + | 1.83 | 1.78E-13 | ncr0570; slr2067 5'UTR, CDS | Ncr0570; allophycocyanin alpha chain ApcA | |
| 37 | 2375275 | 2375808 | + | 1.83 | 3.21E-18 | slr0373 CDS; slr0374 5'UTR, CDS | Ycf46 | both genes are down-regulated at RNA level at cold temperature, more pronounced in WT than in Δ*crhR* (1) |

| | | | | | | | | |
|---|---|---|---|---|---|---|---|---|
| 38 | 958458 | 958918 | + | 1.81 | 1.80E-13 | slr1841 CDS | major outer membrane porin | - |
| 39 | 2082357 | 2082631 | + | 1.80 | 4.66E-13 | slr0442 CDS | hypothetical protein, 'target gene of Sycrp1' | - |
| 40 | 3284260 | 3284583 | - | 1.77 | 9.91E-16 | sll1338 CDS | hypothetical protein | - |
| 41 | 2779346 | 2779673 | + | 1.77 | 6.04E-13 | slr0906 CDS | PSII CP47 reaction center protein PsbB | lower RNA levels in WT and mutant in the cold (1) |
| 42 | 2181021 | 2181417 | + | 1.69 | 8.42E-10 | slr0151 CDS | PSII assembly proteins operon | somewhat lower RNA level in the cold in WT and in Δ*crhR* (1) |
| 43 | 8033 | 8361 | + | 1.66 | 3.81E-14 | slr1311 CDS, 3’UTR | PSII D1 protein PsbA2 | - |
| 44 | 596948 | 597328 | + | 1.65 | 3.50E-14 | slr2051 CDS, 3'UTR | PBS rod-core linker polypeptide CpcG1 | strongly reduced RNA levels in WT and mutant in the cold (1) |
| 45 | 7555 | 7889 | + | 1.63 | 4.26E-12 | slr1311 CDS | PSII D1 protein PsbA2 | - |
| 46 | 983763 | 984020 | + | 1.59 | 1.44E-10 | slr1545 5'UTR, CDS | extracytoplasmic function (ECF) sigma factor SigG | lower RNA level in the cold in WT and in Δ*crhR* (1) |
| 47 | 2526177 | 2526342 | - | 1.54 | 2.15E-12 | sll0199 5'UTR, CDS; ncl1260-ncRNA | plastocyanin PetE; Ncl1260 | - |
| 48 | 596449 | 596764 | + | 1.50 | 1.50E-08 | slr2051 5'UTR, CDS | PBS rod-core linker polypeptide CpcG1 | strongly reduced RNA levels in WT and mutant in the cold (1) |
| 49 | 3457392 | 3457658 | + | 1.46 | 1.91E-07 | slr1655 5'UTR, CDS | PSI subunit XI PsaL | strong down-regulation of RNA level at the lower temperature (1); consistent with lower PSI trimer-to-monomer ratio in the cold |
| 50 | 2377279 | 2377584 | + | 1.42 | 2.97E-08 | slr0376 CDS; ncr1200 | Ycf46 | down-regulated at RNA level at cold temperature, more pronounced in WT than in Δ*crhR* (1) |
| 51 | 2961084 | 2961428 | + | 1.42 | 5.20E-09 | slr0623 5'UTR, CDS | thioredoxin A | RNA level is cold-repressed in WT and in Δ*crhR* (1) |
| 52 | 2612825 | 2613041 | + | 1.38 | 6.91E-08 | slr0476 5'UTR, CDS | hypothetical protein | divergent regulation in WT and Δ*crhR* upon transfer to lower temperature: RNA |

| | | | | | | | | |
|---|---|---|---|---|---|---|---|---|
| | | | | | | | | level down in WT but up in Δ*crhR* (1) |
| 53 | 2644699 | 2645018 | + | 1.34 | 2.15E-08 | slr0628 5'UTR, CDS | 30S ribosomal protein S14, RpsN | consistent with reported effects on ribosomal proteins (2); cold induction seen in WT is almost missing in Δ*crhR* (1) |
| 54 | 957855 | 958275 | + | 1.27 | 1.31E-07 | slr1841 5'UTR, CDS | major outer membrane porin | - |
| 55 | 726665 | 726967 | - | 1.20 | 3.53E-07 | sll1578 CDS; sll1577 CDS | C-phycocyanin alpha chain CpcA; C-phycocyanin beta chain CpcB | cold repression of both genes at RNA (1) and protein level (2), more pronounced in WT than in Δ*crhR*) |
| 56 | 2525814 | 2526138 | - | 1.19 | 1.81E-07 | sll0199 CDS | plastocyanin PetE | - |
| 57 | 7177 | 7515 | + | 1.13 | 5.29E-06 | slr1311 5'UTR, gene | PSII D1 protein PsbA2 | - |
| 58 | 959758 | 960088 | + | 1.08 | 3.25E-04 | slr1841 CDS | major outer membrane porin | - |
| 59 | 2046872 | 2047105 | + | 1.01 | 2.01E-04 | slr1634 5'UTR, gene | hypothetical protein | strongly reduced RNA levels in WT and mutant in the cold (1) |
| 60 | 3284627 | 3284918 | - | 0.91 | 4.92E-04 | sll1338 CDS | hypothetical protein | - |
| 61 | 1982014 | 1982422 | + | 0.89 | 6.97E-04 | ssr2831-5'UTR, CDS; slr1689 CDS | PSI reaction center subunit IV, PsaE; formamidopyrimidine-DNA glycosylase MurM | strongly reduced RNA levels in WT and mutant in the cold (1) |
| 62 | 2791013 | 2791413 | + | 0.87 | 1.59E-04 | 6803t34- exon 1, slr0915-5'UTR, CDS | tRNA(Leu)$^{UAA}$ exon 1, endonuclease slr0915 | - |
| 63 | 2695562 | 2695828 | + | 0.87 | 4.06E-04 | ssr0692-5'UTR, CDS | N-regulatory protein PirA, controls flux into ornithine-ammonia cycle | somewhat upregulated RNA levels in both strains in the cold (1) |
| 64 | 2539363 | 2539697 | - | 0.83 | 1.73E-03 | sll0416 CDS | groEL2, 60 kDa chaperonin 2 | upregulated RNA levels in both strains in the cold (1) |
| 65 | 1819735 | 1820003 | - | 0.75 | 1.65E-03 | sll1867-5'UTR, CDS | PSII protein D1 PsbA3 | - |

| | | | | | | | | |
|---|---|---|---|---|---|---|---|---|
| 66 | 2525624 | 2525787 | - | 0.74 | 4.53E-03 | sll0199-3'UTR | 3'UTR of petE | - |
| 67 | 726355 | 726645 | - | 0.72 | 2.68E-03 | sll1578 CDS | C-phycocyanin alpha chain CpcA | cold repression at RNA (1) and protein level (2), more pronounced in WT than in Δ*crhR* |
| 68 | 2705881 | 2706279 | - | 0.66 | 1.50E-02 | sll0376 CDS; ncl1350-ncRNA | hypothetical protein, Ncl1350 | - |
| 69 | 2471690 | 2471734 | - | 0.61 | 2.02E-02 | sll0018-5'UTR | fructose-bisphosphate aldolase class 2 FbaA | somewhat reduced RNA levels in WT but not in the mutant in the cold (1) |
| 70 | 2468191 | 2468633 | - | 0.60 | 2.43E-02 | sll0020 CDS | ATP-dependent Clp protease regulatory subunit ClpC | - |
| 71 | 1297460 | 1297562 | - | 0.60 | 2.08E-02 | sll1694-5'UTR, CDS | type 4 pilin-like protein PilA1 | down-regulation in the cold in both strains at RNA level (1) |
| 72 | 2484894 | 2485282 | - | 0.56 | 3.50E-02 | ssl0020-5'UTR, CDS | ferredoxin I, PetF | slight down-regulation in the cold at RNA level (1) |
| 73 | 1296934 | 1297338 | - | 0.53 | 2.45E-02 | sll1694 CDS | type 4 pilin-like protein PilA1 | down-regulation in the cold in both strains at RNA level (1) |
| 74 | 727336 | 727720 | - | 0.50 | 4.09E-02 | sll1577-5'UTR, CDS; ncr0290-ncRNA | C-phycocyanin beta chain CpcB; Ncr0290 | cold repression of *cpcB* and following gene *cpcA* at RNA (1) and protein level (2), more pronounced in WT than in Δ*crhR* |
| 75 | 724505 | 724971 | - | 0.48 | 4.68E-02 | sll1580 CDS | PBS 32.1 kDa linker polypeptide CpcC1 | cold induction observed at protein level in WT (2) but repression at RNA level in Δ*crhR* and, more pronounced, in WT (1); this points at divergent regulation caused by CrhR |

### Supplementary Figures

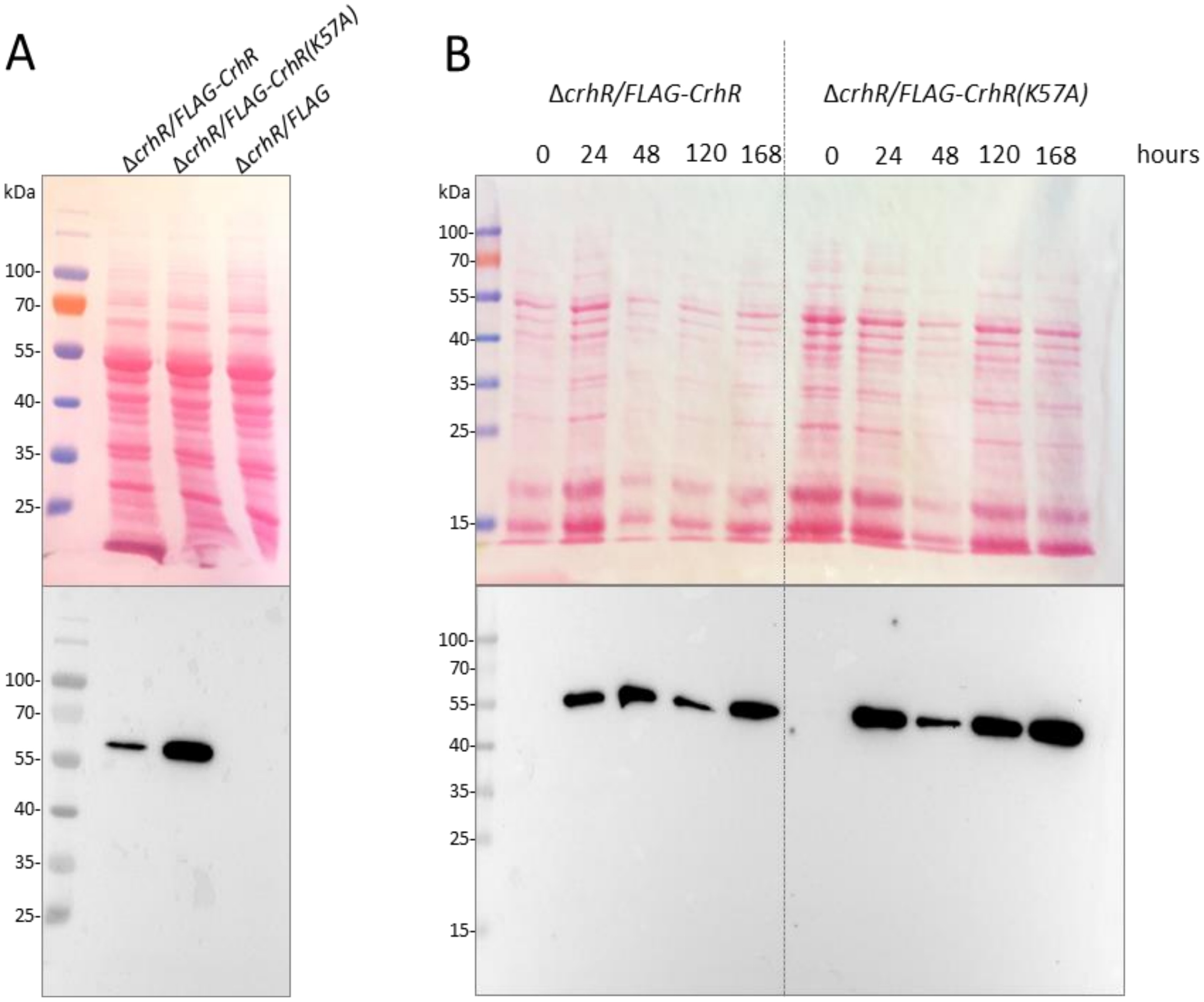


**Fig. S1. Western blot analysis for the detection of FLAG-tagged CrhR proteins in different strains of *Synechocystis*.** Soluble proteins isolated from cells grown at 30°C (10 µg) were separated by electrophoresis on 10% SDS-polyacrylamide gels and transferred to nitrocellulose membranes. The blots were stained with Ponceau S solution to control for loading (top) and probed with ANTI-FLAG® M2-Peroxidase (HRP) antibody (bottom). **(A)** Lysate fraction containing soluble proteins isolated 24 hours after induction with 2 µM $CuSO_4$ from $CrhR_{WT}$, $CrhR_{K57A}$ and $CrhR_{control}$ strains. **(B)** Soluble proteins from $CrhR_{WT}$ and $CrhR_{K57A}$ strains. Aliquots were harvested for protein isolation before the induction by addition of $CuSO_4$ (time point 0) and 24, 48, 120, 168 hours after the induction with 2 µM $CuSO_4$. Strains were grown in Erlenmeyer flasks in BG-11 medium.

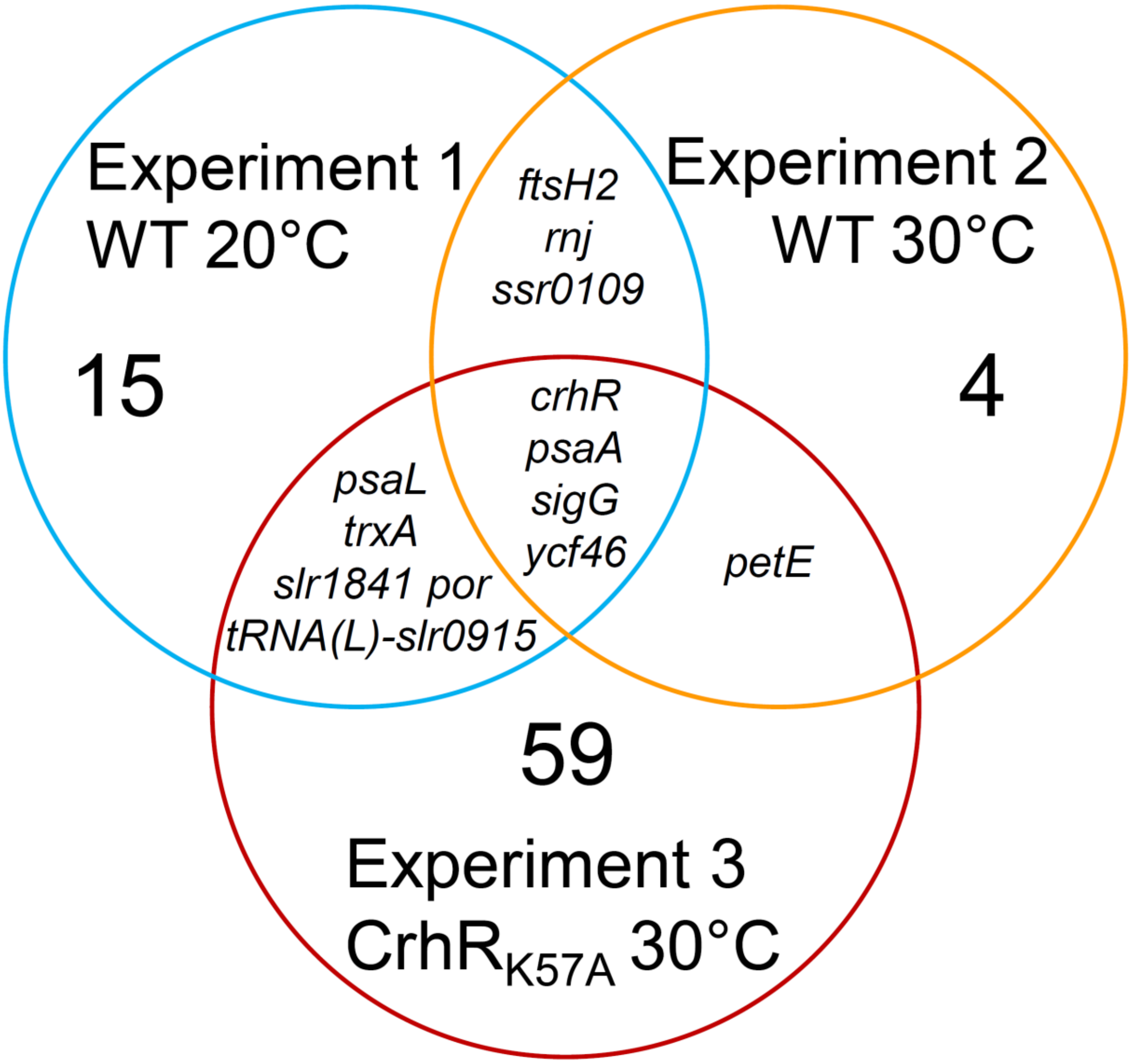


**Fig. S2. Overlaps in the identified transcripts between the three performed experiments.** In total, 119 peaks were recovered belonging to 90 different genes. If more than a single peak was recovered for a gene, these peaks were counted only once in the here depicted diagram. Further details, including the exact coordinates of recovered transcripts, enrichment factors and adjusted p values in each of the experiments, are given in **Tables S3 to S5**.

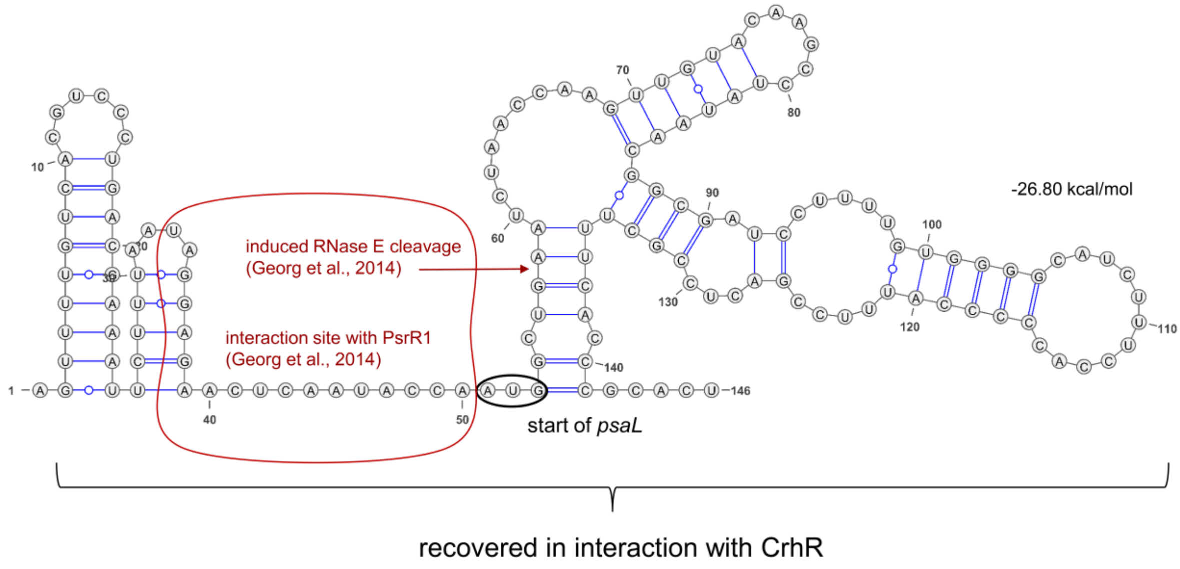


**Fig. S3. Predicted secondary structures in mRNA segments encompassing the 5' UTR and the initial ORF of *psaL* that were recovered in CrhR co-IP experiments.** The interaction site relevant for the PsrR1-mediated post-transcriptional downregulation of *psaL* translation and mRNA destabilization by recruiting RNase E after shift to high light (Georg *et al.*, 2014) are indicated.
